# Increased TCR signaling in regulatory T cells is disengaged from TCR affinity

**DOI:** 10.1101/2023.01.17.523999

**Authors:** Yi Jing, Yuelin Kong, Denise Allard, Baoyu Liu, Elizabeth Kolawole, Maran Sprouse, Brian Evavold, Matthew Bettini, Maria Bettini

## Abstract

Foxp3+ regulatory T cells (Tregs) are capable suppressors of aberrant self-reactivity. However, TCR affinity and specificities that support Treg function, and how these compare to autoimmune T cells remain unresolved. In this study, we used antigen agnostic and epitope-focused analyses to compare TCR repertoires of regulatory and effector T cells that spontaneously infiltrate pancreatic islets of non-obese diabetic mice. We show that effector and regulatory T cell-derived TCRs possess similar wide-ranging reactivity for self-antigen. Treg-derived TCRs varied in their capacity to confer optimal protective function, and Treg suppressive capacity was in part determined by effector TCR affinity. Interestingly, when expressing the same TCR, Tregs showed higher Nur77-GFP expression than Teffs, suggesting Treg-intrinsic ability to compete for antigen. Our findings provide a new insight into TCR-dependent and independent mechanisms that regulate Treg function and indicate a TCR-intrinsic insufficiency in tissue-specific Tregs that may contribute to the pathogenesis of type 1 diabetes.

## Introduction

Type 1 diabetes (T1D) is characterized by autoimmune targeting of beta cells, which is mediated by anti-beta cell antigen T cell responses^1^. CD4+Foxp3+ Regulatory T cells (Tregs), however, are effective at suppression of pancreatic autoimmunity, but ultimately fail in T1D^2^. The exact antigens targeted by autoimmune vs regulatory T cells in T1D, and differences in T cell receptor biophysical parameters that support Treg vs effector T cell function in autoimmunity are not fully resolved. The thymic development of Tregs is directed by TCR specificity and affinity and is heavily dependent on the thymic presentation of cognate antigens^3-6^. As a result, tTregs express a diverse TCR repertoire that has strong self-reactivity and minimal overlap with Foxp3-effector T cells (Teffs)^7-9^. However, the reported lack of TCR overlap and uniqueness of the Treg TCR repertoire is primarily derived from sequencing polyclonal T cells without focusing on a particular antigenic specificity. Indeed, when a single antigen specificity is explored, the level of repertoire overlap between Treg and Teff can be as high as 40%^10,11^. Moreover, in experimental autoimmune encephalomyelitis (EAE) model for multiple sclerosis, Treg and Teff TCRs were shown to exhibit similar range of affinity and reactivity for myelin oligodendrocyte glycoprotein^12^. These observations suggest that high self-reactivity does not guarantee development into a Treg lineage, and low affinity for self-antigen in some cases might be sufficient for Treg development. However, due to the rarity of Treg cells of a single peptide specificity, it remains unclear whether Treg development and recruitment to the tissue in a polyclonal environment can be supported by a wide range of TCR reactivities, and whether all tissue Treg-derived TCRs are functional in protection against autoimmunity.

Tissue specificity is thought to be necessary for optimal Treg efficacy in T1D, but our limited understanding of how TCR specificity and affinity control Treg function is an obstacle to developing optimal antigen specific Treg therapies^13-15^. In our previous studies we observed that Treg function in non-obese diabetic (NOD) mouse model positively correlated with high Treg self-reactivity, as defined by TCR signaling (Nur77-GFP) and CD5 expression^16^. However, Tregs expressing a low insulin affinity TCR were also capable of contributing to regulation of autoimmune diabetes, while exhibiting distinct transcriptional profile^16,17^, indicating that high and low affinity Tregs may play non-redundant roles in T1D prevention. On the other hand, Treg function is less sensitive to affinity increase through TCR engineering compared to Teffs^18^.

In this study, we utilized antigen agnostic and insulin-tetramer specific T cell isolation to compare TCR repertoires of pancreatic islet infiltrating Treg and Teff cells in the context of autoimmune diabetes. Both approaches revealed a similar breadth of reactivities to islet antigens between Tregs and Teffs, despite higher Nur77-GFP and CD5 expression in islet-infiltrating Tregs. Using TCR retrogenic mice, adoptive transfer, and TCR-redirected Tregs we show that Treg-derived TCRs vary in their ability to confer optimal Treg function. Interestingly, Tregs exhibited a cell-intrinsic capacity to translate signals through low affinity TCRs into increased downstream signaling compared to Teffs; however, this increase was not sufficient to overcome insufficient TCR function under inflammatory stress. In summary, we detected a TCR-intrinsic insufficiency in tissue-specific Tregs that may contribute to the pathogenesis of T1D.

## Results

### Tissue-infiltrating Tregs in autoimmunity have low reactivity for self-antigen

To determine antigen specificities and relative antigen reactivity of islet infiltrating CD4+ effector and Foxp3+ regulatory T cells, 14 TCRs (7 from Teffs and 7 from Tregs) were cloned directly from infiltrated islets of a prediabetic NOD.Foxp3-GFP female mouse (Table 1). T cells expressing isolated TCRs were generated using TCR retrogenic approach (TCR Rg) as previously described^19,20^. T cells isolated from spleens of single TCR Rg mice were expanded *in vitro* and stimulated with the two dominant CD4 epitopes targeted in NOD diabetes: insulin B:9-23 and chromogranin/insulin hybrid peptide 2.5HIP^21-23^. Reactivity to insulin was common in both Teff-(4/7) and Treg-derived TCRs (3/7), and one Teff TCR was reactive to 2.5HIP (TE-F7) (Table 1, Figure S1), confirming previous studies that insulin reactivity is dominant among islet infiltrating CD4+ T cells^24-26^. Based on previously documented high self-reactivity of islet-infiltrating Tregs as evidenced by higher levels of CD5 and Nur77 compared to Teffs^16,27^, we expected to see increased reactivity of Treg-derived TCRs to the insulin epitope. Contrary to our expectations, both populations showed a similar range of IFNγ response (0.5% to 4%) to insulin peptide (Figure S1).

**Table 1.**
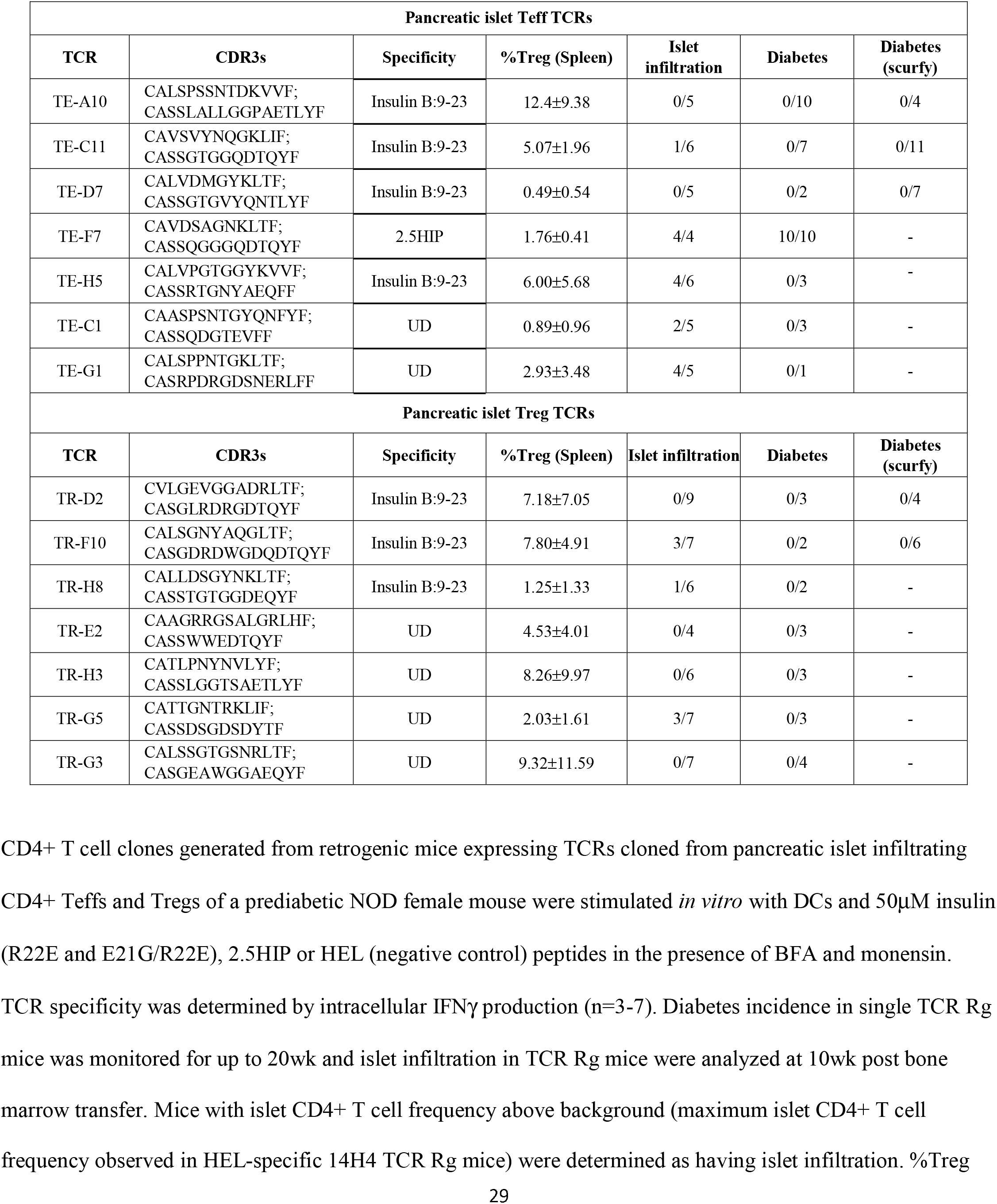
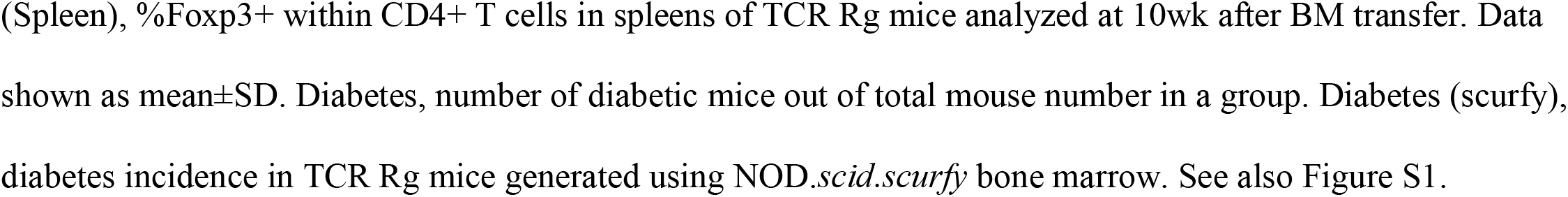
Summary of Treg and Teff TCRs isolated from pancreatic islets of a prediabetic NOD female.

When islet-infiltrating cells in NOD mice were stained with insulin B:10-23(E21G/R22E) tetramer (Ins-tet) (Figure 1A), Tregs exhibited higher frequencies of Ins-tet+ cells (Figure 1A&1B). Importantly, after gating on Ins-tet+ cells, Tregs still showed significantly higher expression levels of CD5 and Nur77-GFP (Figure 1C&1D), indicating that the increased self-reactivity in Tregs was not dependent on different TCR specificities compared with Teffs. Therefore, we considered the possibility that Treg TCRs recognized less dominant epitopes besides the well-characterized InsB:9-23 and 2.5HIP *in vivo*. To address this possibility, we monitored Treg and Teff TCR Rg mice for T cell infiltration into the pancreas and diabetes development. As summarized in Table 1, at 10 weeks post bone marrow (BM) transfer, 8/14 TCRs showed infiltration above background established using a control hen egg lysozyme specific TCR. However, no spontaneous diabetes development was observed except for the 2.5HIP reactive TE-F7. Since the majority of TCRs supported Treg development, we confirmed that endogenous Tregs did not restrain autoimmunity by selecting 5 insulin reactive TCRs (3 Teff and 2 Treg) to be expressed using NOD.*scid*.*scurfy* (Foxp3 mutant) BM that prevents development of functional Tregs^28,29^. Again, diabetes did not develop even in the absence of functional Tregs (Table 1).

**Figure 1.**
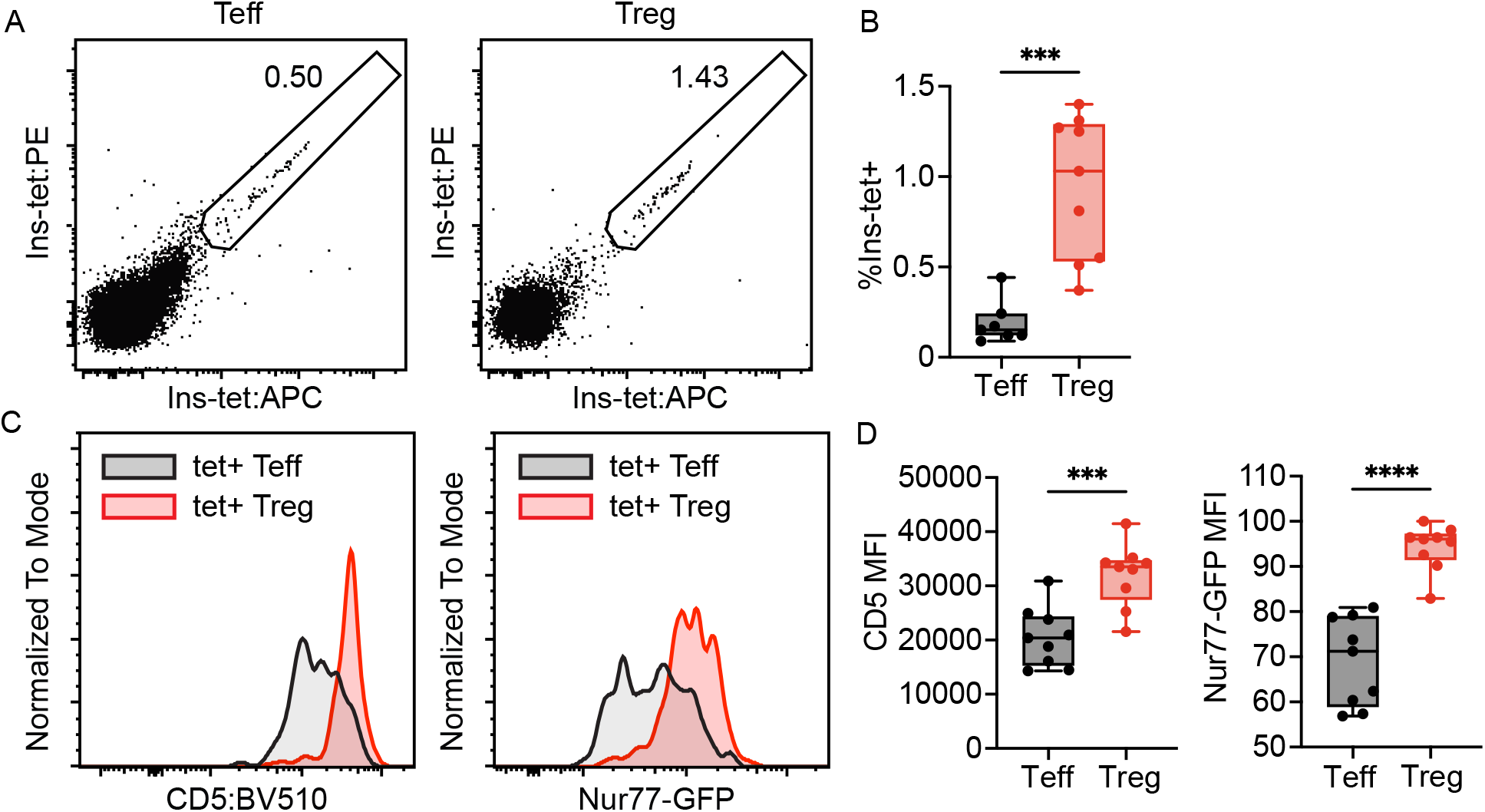
Islet-infiltrating insulin specific Tregs showed higher level of self-reactivity and TCR signaling than Teffs. Teffs and Tregs were isolated from infiltrated pancreatic islets of pre-diabetic NOD.Nur77-GFP females and were stained with insulin B:10-23(E21G/R22E) tetramer. **(A, B)** Representative flow plots and summary of Ins-tet+ frequencies in islet Teffs and Tregs. **(C, D)** Representative flow plots and summary of CD5 and Nur77-GFP expression levels. Data were analyzed by (B) paired t test and (D) Wilcoxon test (n=9). Outliers were identified using the ROUT method (Q=1) and removed from analysis. ****, p ≤ 0.0001; ***, p ≤ 0.001; **, p ≤ 0.01; *, p ≤ 0.05; ns > 0.05.

Our observations suggest a general lack of diabetogenicity among islet-specific TCRs, which is consistent with an idea that diabetes is driven by few initiator clones that preferentially recognize islet neoantigens not expressed in the thymus, and therefore lack tTreg development of the same specificity^30-32^. However, if islet antigen specific tTregs are skewed towards increased affinity for antigens expressed in the thymus, such as insulin, we would expect Treg repertoire to be enriched in TCRs with high capacity for beta cell destruction, which we did not observe. Our unexpected observations that Tregs possess TCRs with similar reactivity for insulin self-antigen as Teffs prompted us to perform a broad TCR repertoire analysis.

### TCR repertoire of islet-infiltrating effector and regulatory T cells

We conducted scTCRαβ analysis of CD4+ T cells isolated from pancreatic islets of 4 prediabetic NOD females (Figure S2A-S2C). Consistent with previous studies, analysis of CD4+Foxp3-Teffs and Foxp3+ Tregs revealed the uniqueness of islet Treg repertoire with a low level of repertoire overlap (0.62%±0.32%) and similarity measured by Morisita index (Figure 2A&2B, Figure S2D). Repertoire diversity was significantly higher in Teffs (Figure 2C), while repertoire evenness, measured by the inverse Pielou index was not significantly different (Figure 2D). Consistently, top 10 clones made up 10% and 17% of Teff and Treg repertoires, respectively (Figure 2E), suggesting similar ability of Treg and Teff clones to expand in the tissue. Length of both CDR3α and CDR3β fit Poisson distribution with means of 13.5 and 14 amino acids respectively (Figure S2E). While no bias in TRAV gene usage was observed between Tregs and Teffs (Figure S2F), TRBV10 and TRBV15 were overrepresented in Teffs and TRBV12-1 in Tregs (Figure 2G). Interestingly, the most prevalent V-alpha and V-beta segments (TRAV5-4, TRAV6N7; TRBV5; TRBV12-2+TRBV13-2) lost their dominance when we focused our analysis on clonotypes shared between Tregs and Teffs (Figure S2G), suggesting that their prevalence was not a result of expansion of converted clones.

**Figure 2.**
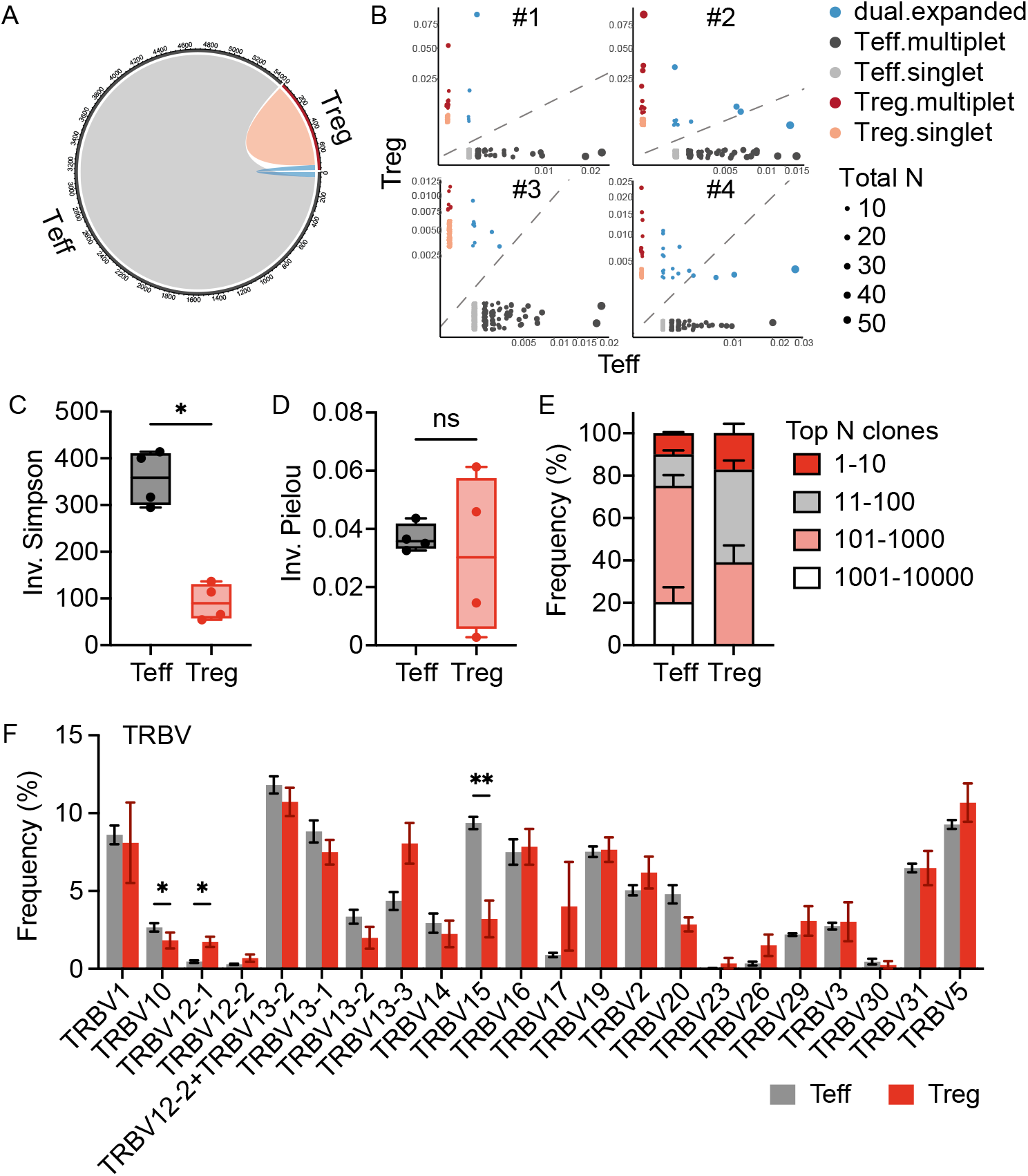
TCR repertoire of pancreatic islet infiltrating CD4+ Treg and Teff cells. Single cell RNAseq analysis of TCRaβ repertoires of Teffs and Tregs isolated from pancreatic islets of four 12-week-old pre-diabetic NOD females. **(A, B)** TCR repertoire overlap between Treg and Teff cells (A) pooled or (B) individually shown. Dot size represents numbers of cells included in a clonotype (CDR3 amino acids). Frequencies (0-1) within Teff and Treg repertoires were plotted on x- and y-axis, respectively. **(C)** Inverse Simpson and **(D)** inverse Pielou index of islet Tregs and Teffs TCR repertoires. **(E)** Clonal expansion level of islet Tregs and Teffs. **(F)** TRBV gene usage. Data are plotted as mean±SEM (n=4). **, p ≤ 0.01; *, p ≤ 0.05; ns > 0.05 by (C, D) Mann-Whitney test and (F) multiple paired t test. See also Figure S2.

### Insulin specific Tregs and Teff exhibit similar range of TCR affinities

To compare TCR repertoires of insulin specific Tregs and Teffs, we generated TCR Rg mice that expressed an endogenous polyclonal TCRβ repertoire and a fixed TCRα cloned from a high affinity insulin B:9-23 reactive TCR P2 (P2-TCRα mice) (Figure 3A)^28^. Although Ins-tet binding cells were extremely rare in the spleen and not enriched over CLIP tetramer binding cells, their frequency significant increased in pancreatic islet, enabling downstream analysis (Figure S3A&S3B). Islet Ins-tet+ Tregs and Teffs were compared as in Figure 1, confirming a significantly higher expression level of CD5 on Ins-tet+ Tregs (Figure 3B&3C). To compare Treg and Teff TCR affinities restricted by the same epitope, we applied 2D micropipette adhesion assay^33^. We sorted Teffs (Foxp3-GPF-) and Tregs (Foxp3-GFP+) from pancreatic islets of P2-TCRα mice and tested T cell binding to insulin B:9-23(E21G/R22E):I-A^g7^ complex presented on human red blood cell attached to a micropipette. Based on positive adhesion frequencies, 70% Teffs and 100% Tregs were determined to be insulin specific (Figure 3D). Interestingly, we also observed no significant difference in TCR affinity for insulin epitope between Tregs and Teffs (Figure 3E), with both populations showing a range of affinities spanning at least 100-fold.

**Figure 3.**
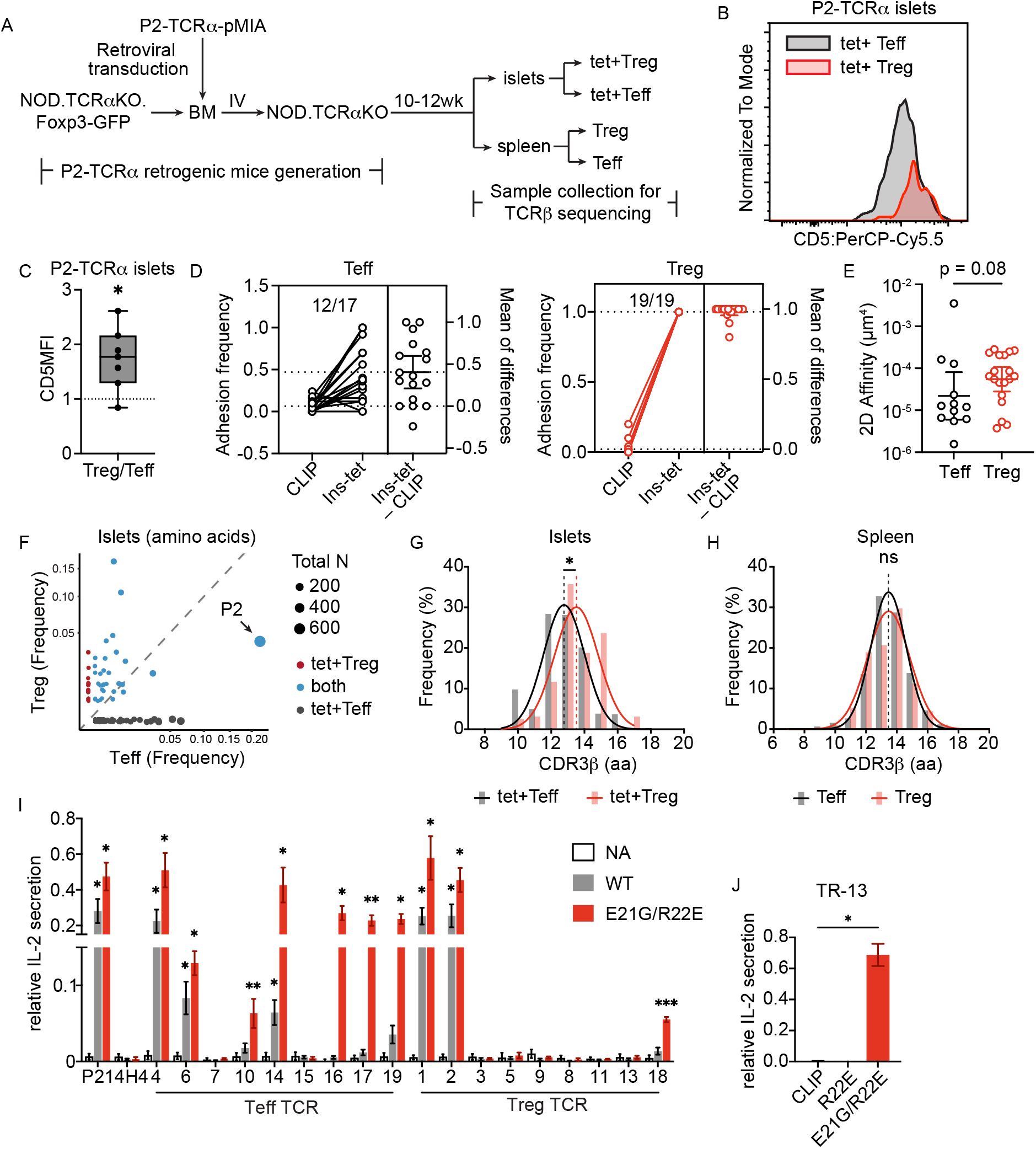
Pancreatic islet-infiltrating tetramer positive Tregs exhibit a wide range of reactivity to insulin and possess unique TCR characteristics compared with Teffs. **(A-H)** Bone marrow (BM) hematopoietic stem cells collected from NOD.TCRαKO.Foxp3-GFP mice were transduced with a retroviral vector that encodes the TCRα chain of insulin specific TCR P2 and an Ametrine expression marker. Transduced BM cells were transferred intravenously into irradiated NOD.TCRαKO recipients. Recipients were sacrificed 10-12wk later and Ametrine+CD4+ T cells were isolated from spleens and pancreatic islets. Splenic GFP+ Tregs and GFP-Teffs were sorted individually. Islet samples from 38 mice were pooled and stained with Ins-tet. Ins-tet+ Teffs and Tregs were sorted. Genomic DNA was purified from sorted samples and TCRβ repertoires were sequenced. **(A)** Diagram for experimental process. **(B&C)** CD5 expression in Ins-tet+ Tregs and Teffs isolated from pancreatic islets of P2-TCR α mice. Data are pooled from 3 independent experiments (n=7) and analyzed by one sample t test. **(D&E)** Eight pre-diabetic P2-TCRα females were sacrificed 14 weeks after BM transfer. Teffs and Tregs were sorted from pancreatic islets. Specificity of sorted cells were measured using the 2D micropipette adhesion frequency assay against insulin B:10-23(E21G/R22E) and CLIP in the context of I-A^g7^. (D) T cell-pMHC adhesion frequencies. Dash lines, population means. (E) 2D affinity to insulin. Data are plotted as geometric means ± 95%CI and analyzed by Mann-Whitney test. **(F)** TCR repertoire (amino acid) overlap between islet tet-Ins+ Tregs and Teffs. Dot size represents numbers of cells included in a specific clonotype. The dominant T cell clone which expressed the same TCRβ as TCR P2 was labeled. (G&H) Distributions of CDR3β length in **(G)** islet Ins-tet+ and **(H)** splenic repertoires. Extra sum-of-squares F test. **(I&J)** Sequencing identified TCRs were expressed in 4G4 thymoma cells. TCR+4G4 cells were stimulated with (I) 10μM of wild-type insulin B:9-23 peptide (WT) or insulin B:9-23 (E21G/R22E) agonist mimotope and M12.C3.g7 cells or (J) plate-bound insulin or control CLIP tetramers for 24hr. IL-2 concentrations in the supernatant were measured by ELISA and normalized to antiCD3 positive control. Data are plotted as means ± SEM and are (I) pooled from or (J) representative of 3 independent experiments and analyzed by Welch ANOVA followed by Benjamini, Krieger and Yekutieli’s test. All comparisons are made with negative control. ***, p ≤ 0.001; **, p ≤ 0.01; *, p ≤ 0.05; ns > 0.05. See also Figure S3-S5, Table S1, S2.

For TCR sequencing, Foxp3-GFP reporter and tetramer staining were used to ensure purity of isolated insulin specific Teffs and Tregs from pancreatic islets of P2-TCRα mice. TCRβ sequences were obtained from 3837 Ins-tet+ Teffs and 499 Tregs isolated from combined islets of 38 P2-TCRα mice. TCRβ repertoires of splenic Tregs and Teffs of 5 individual mice were simultaneously sequenced without pooling or tetramer staining and were used as a reference library. A total of 358 unique Teff clonotypes and 92 unique Treg clonotypes at amino acid level were obtained from the islets (Table S1), in which 35 were shared between Tregs and Teffs (Table S1, Figure 3F). Interestingly, this relatively high level of repertoire similarity was also observed in the spleens (Table S1), suggesting an effect of the restricted TCR repertoire rather than iTreg conversion in the tissue. TRBV13-02 usage was identified most frequently for both Tregs and Teffs, followed by TRBV02-01 and TRBV31-01. While the preference for TRBV13-02 was also seen in the spleen and was likely because that P2 TCRα originally paired with TRBV13-02, preference for TRBV02-01 and TRBV31-01 was limited to islets (Figure S3C&S3D). Compared with Tregs, islet Ins-tet+ Teffs tended to express shorter CDR3β, which was not observed in splenic repertoires (Figure 3G&3H). Overall, our data suggest that islet-infiltrating insulin specific Tregs and Teffs preferred TCRs with distinct characteristics.

### Thymic antigen encounter imprints on islet TCR repertoires

To confirm that insulin-specific Treg TCR repertoire characteristics described above were dependent on thymic selection and expression of insulin, we utilized NOD.Ins1KO.Ins2KO.InsY16A.TCRαKO (NOD.Ins^Y16A^.TCRαKO) mice, which express a form of insulin that is mutated at the key TCR contact residue (Y16A) that disrupts T cell recognition^6,34^. As diagramed in Figure S4A, we generated P2-TCRα mice using NOD.Ins^Y16A^.TCRαKO as BM recipients (P2-TCRα.Y16A mice). CD4+ T cells were isolated from the spleens of P2-TCRα.Y16A mice and transferred into NOD.TCRαKO recipients to allow T cell recognition of insulin and subsequent recruitment to the pancreas. Compared with P2-TCRα mice generated under conditions of normal insulin expression, a significant decrease in islet Ins-tet+ Tregs was observed in mice that received P2-TCRα.Y16A-derived splenic CD4+ T cells, suggesting that peripheral conversion, if any, was not sufficient to compensate for the reduced tTreg development (Figure S4B&S4C). Islet samples were collected from 25 T cell recipients generated from 7 P2-TCRα.Y16A donors, pooled, and a total of 9544 Teff and 146 Treg TCRβ chains were obtained through sequencing, which included 210 unique Teff and 35 Treg clonotypes (Table S1).

Ten clonotypes were shared between Teffs and Tregs (4.2% of all clonotypes, 28.6% of Tregs; Figure S4D). The preference for shorter CDR3βs was no longer observed in Tregs (Figure S4E). Instead, both Tregs and Teffs strongly favored CDR3βs with 13 amino acids. TRBV13-02, TRBV02-01 and TRBV31-01 remained the three most dominant TRBV gene identified in Teffs; however, TRBV02-01 and TRBV31-01 lost their dominance in Tregs (Figure S4F), supporting a role for thymic insulin in shaping Treg TCR repertoire. Together, our data demonstrate nonredundant imprints by thymic antigen encounter and TCR self-antigen specificity on islet Treg TCR repertoires.

### Treg TCRs selected from P2-TCRα islet repertoire exhibit a broad range of insulin reactivity

To compare Treg and Teff TCR parameters restricted by the same epitope, we selected 18 TCRs from the islet Ins-tet+ repertoires of P2-TCRα retrogenic mice for functional analysis (Table S2) among which 9 were expressed exclusively by Teff cells (henceforth called Teff TCRs), and 9 were expressed by islet Tregs (Treg TCRs). TCR transfectant cell lines were stimulated with insulin peptides and M12.C3.g7 APCs. IL-2 secretion was used as a readout of TCR reactivity. TCRs without significant response to insulin peptides were also stimulated with plate-bound tetramers, which offers better sensitivity. Surprisingly, despite being isolated from Ins-tet+ islet-infiltrating T cells, only 4 out of 9 Treg TCRs showed a significant response to insulin, while 7 of 9 Teff TCRs were confirmed to be insulin specific (Figure 3I&3J). Given that the majority of islet T cells in P2-TCRα mice were insulin specific (Figure 3D), non-responders likely had functional avidity below the threshold of detection. Among responders, both the highest and the lowest reactivity were identified from Treg TCRs, suggesting a broad range of reactivity. At the same time, as observed with TCRs selected from WT NOD islets (Figure S1), Ins-tet+ Treg TCRs did not show an overall higher reactivity over Teff TCRs (Figure 3I&3J, Figure S5). Overall, our observations in NOD and Ins-tet selected TCRs suggest that a large proportion of insulin specific Tregs recruited to the tissue during autoimmunity express low affinity TCRs.

### Treg-derived insulin specific TCRs do not increase Treg development

We then considered the possibility that Treg-derived TCRs are cross-reactive to other self-antigens or recognize a modified version of the insulin epitope unique to the *in vivo* environment. To compare the *in vivo* function of Treg and Teff TCRs, we re-expressed insulin-responsive TCRs selected from Table S2 *in vivo* to monitor T cell recruitment to the tissue and induction of diabetes. Individual TCRs were expressed by transducing NOD.*scid* BM that was then transferred into NOD.TCRαKO recipients. No significant difference was observed in splenic CD4+ T cell number or frequency (Figure 4A, Figure S6A), indicating comparable peripheral engraftment and T cell development. Spontaneous diabetes development was observed in mice expressing both Treg (TR-1, 2, 13) and Teff TCRs (TE-6), and both high (TR-1, TR-2) and low reactivity (TR-13) TCRs (Figure 4B). All except for TE-16 and TR-18 showed significant islet-infiltration compared with the hen egg lysozyme (HEL) specific TCR 14H4 (Figure 4C, Figure S6B). These observations were consistent with our previous findings that TCR reactivity to islet antigen by itself does not predict TCR diabetogenicity^28^, but also showed that Treg TCRs are not uniquely pathogenic due to increase *in vivo* antigen sensing or cross-reactivity.

**Figure 4.**
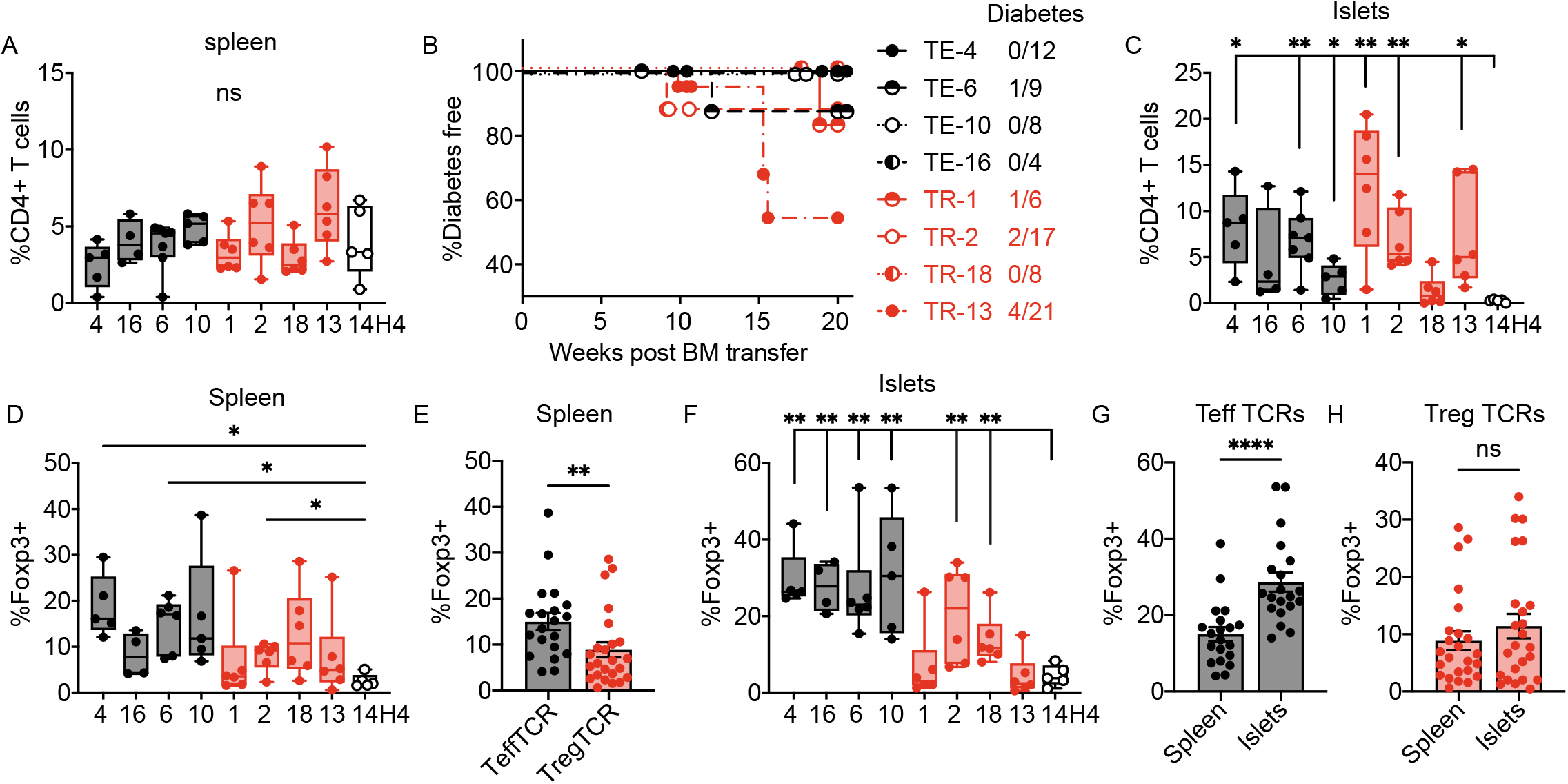
Diabetes and Treg development in Treg and Teff TCR expressing retrogenic mice. Retrogenic mice expressing Teff (black), Treg (red) derived TCRs, or HEL specific 14H4 control TCR (white) were generated by transferring transduced NOD.scid BM into irradiated NOD.TCRαKO recipients. Recipient mice were monitored for diabetes and sacrificed for analysis at diabetes onset or 20wk after BM transfer. **(A)** Frequencies of CD4+ T cells in spleens. **(B)** Frequency of diabetes incidence and diabetic incidence/start-point at-risk mice expressing indicated TCRs. **(C)** Frequencies of CD4+ T cells in pancreatic islets. **(D)** Frequencies of Foxp3+ Tregs among CD4+ T cells in spleens. **(E)** Splenic Treg frequencies with Treg and Teff TCRs analyzed collectively. **(F)** Frequencies of Tregs in pancreatic islets. **(G&H)** Treg frequencies in spleen versus islets of TCR Rg mice expressing (G) Teff- or (H) Treg-derived TCRs. Data were pooled from two independent experiments. ****, p ≤ 0.0001; ***, p ≤ 0.001; **, p ≤ 0.01; *, p ≤ 0.05; ns > 0.05 by (A, C, D, F) One-way ANOVA followed by Benjamini, Krieger and Yekutieli’s test (all comparisons made with 14H4 negative control) and (E, G, H) Mann-Whitney test. See also Figure S6.

Interestingly, robust Treg development was observed in mice expressing Treg TCRs as well as Teff TCRs (Figure 4D). In fact, when TCRs were analyzed collectively, splenic Treg frequency was significantly higher in Teff TCR Rg mice (Figure 4E). Importantly, while TR-1 and TR-2 had the highest reactivity to insulin (Figure 3I), they failed to support better Treg generation than either Treg or Teff TCRs with lower insulin reactivity (Figure 4D). To exclude the possibility Treg and Teff TCR expressing cells had different rate of clonal expansion, we compared splenic Treg frequencies of TCR Rg mice between 10 and 20wk after BM transfer and observed no significant difference (Figure S6C). These observations suggested that the difference between Teff and Treg TCR Foxp3+ frequencies was driven mainly by thymic development and was unlikely a result of peripheral conversion. It has been reported that thymic Treg development is niche sensitive and is limited to a small clonal size^35,36^. To determine whether the lower Treg generation in Treg TCR retrogenic mice was a result of intraclonal competition in the thymus, we co-transferred TCR transduced NOD.*scid* BM with NOD.CD45.2 congenic BM. No significant increase in Treg frequencies or negative correlation between percentages of transduced CD4SP thymocytes and Treg frequencies was observed (Figure S6D&S6E). Our inability to increase Treg development by constraining the niche may reflect a relatively unsaturated status of the insulin-specific T cell niche. This is potentially due to systemic circulation of insulin, and presentation of insulin peptides by APCs in distal lymph nodes^37^. Peripheral circulating DCs contribute to thymic antigen presentation, and could increase the niche for insulin Treg development^37-40^.

The lower frequencies of Tregs in Treg-TCR Rg mice was reflected in the pancreatic islets (Figure 4F). Moreover, while in Teff TCR Rg mice frequencies of Tregs were increased in the islets compared to spleens, such an increase was rarely observed with Treg TCRs (Figure 4G, H). This suggested that in addition to the reduced Treg development, insulin specific Treg TCRs might also be inferior in terms of imparting functional fitness to Tregs. Overall, our data show that for TCRs selected from Ins-tet+ islet repertoires, Treg TCRs did not possess an advantage in supporting Treg development, further demonstrating potential deficiency in antigen recognition by Tregs recruited to the autoimmune response.

### Tregs possess a TCR-independent increased antigen recognition capacity compared to Teffs that express the same TCR

We were puzzled by increased Treg tetramer staining, but similar (or reduced) TCR reactivity of Treg-derived TCRs. We sought to confirm that cloned TCRs still bound the tetramer by labeling Rg single TCR splenocytes with Ins-tet. Unexpectedly, Ins-tet staining was higher on Tregs than Teffs that expressed the same TCR (Figure 5A&5B). This was observed with multiple TCRs, and was not due to different TCR expression level, as we gated on a similar narrow range of CD3 (Figure S7A&S7B). Interestingly, PD-1 was expressed at a higher level by tetramer negative cells, implying biological difference between high and low tetramer binders (Figure S7C). PD-1 colocalizes with TCR in micro-clusters and has been reported to inhibit TCR synapse formation and TCR-pMHC interaction in a PDL1 independent manner^41-43^. Whether PD-1 can regulate T cell antigen recognition potential in the absence of antigen presenting cells or if other negative regulatory mechanisms associated with PD-1+ phenotype can influence TCR/pMHC avidity interactions needs further investigation.

**Figure 5.**
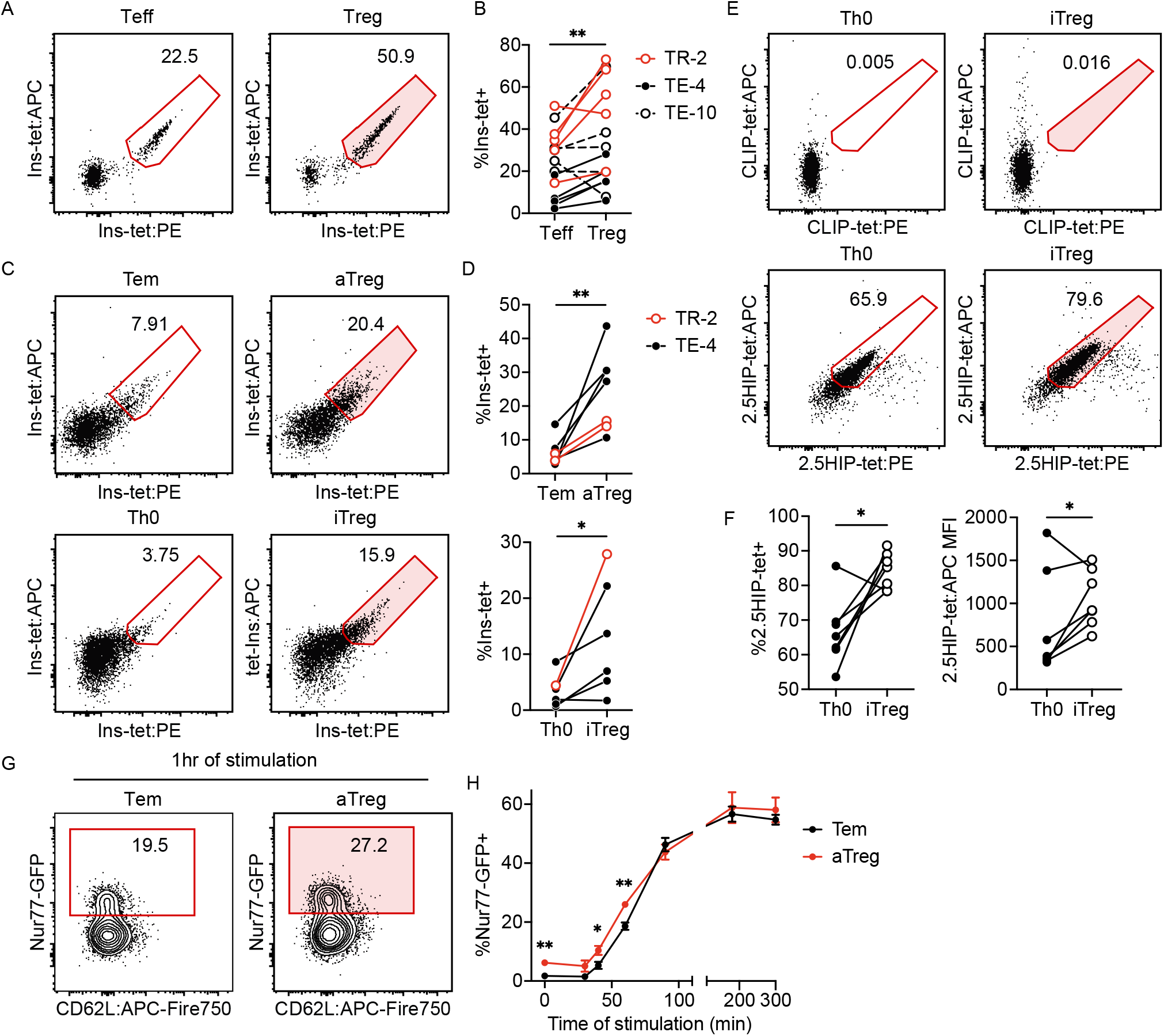
Tetramer staining and stimulation revealed Treg-intrinsic increase in antigen recognition capacity. **(A&B)** Splenocytes of TCR Rg mice were stained with Ins-tet (20min RT followed by 40min on ice). Frequencies of Ins-tet+ cells of Tregs and Teffs with comparable surface CD3 expression were compared. **(C&D)** Tem, aTreg and Tnaive cells were isolated from spleen and ndLNs of NOD mice and transduced with indicated TCRs in vitro. Tnaive cells were differentiated under Th0 and iTreg conditions. Tem, aTreg, Th0 and iTreg cells were stained with Ins-tet for 1hr on ice. Tetramer staining on cells with similar Ametrine expression was compared. **(E&F)** TCR Vβ 4+ Tnaive cells were sorted from spleens of BDC2.5 transgenic mice. Tnaive cells were differentiated under Th0 and iTreg conditions followed by 2.5HIP and CLIP tetramer staining for 1h on ice. (E) Representative flow plots. (F) Summary of tetramer staining. **(G&H)** TE-4 transduced aTregs and Tem cells were stimulated in vitro with plate-bound Ins-tet. Nur77-GFP expression in cell stimulated with plate-bound BDC2.5 tetramer for 5h were used as the zero time-point. Data obtained from the same mouse (B, n=15; F, n=5) or experiment (D, n=6; H, n=4) were aligned and analyzed by paired t test or two-way ANOVA with repeated measures followed by Benjamini, Krieger and Yekutieli’s test. **, p ≤ 0.01; *, p ≤ 0.05; ns > 0.05. See also Figure S7.

To test whether the difference in tetramer binding was a result of different activation status between Tregs and Teffs, we sorted polyclonal activated Tregs (CD44+CD62L-CD25+GITRhi, aTregs) and effector memory T cells (CD62L-CD44+CD62L-CD25-GITRlo, Tem cells) from spleens and non-pancreas draining lymph nodes (ndLNs) from NOD mice. Sorted Tem cells and aTregs were transduced with an insulin specific TCR and stained with Ins-tet on ice to prevent tetramer-induced activation or TCR internalization. Level of tetramer staining positively correlated with expression of the fluorescent reporter of TCR (Ametrine MFI, Figure S7D&S7E). Interestingly, aTregs showed significantly higher Ins-tet staining than Tem cells after gating on a similar narrow level of Ametrine expression (Figure 5C&5D). To test whether the increased Treg capacity for tetramer binding is imprinted on Tregs through their *in vivo* activation or function, or is Treg lineage dependent, we induced Treg differentiation *in vitro* and again observed a higher level of tetramer staining in insulin-TCR transduced iTregs compared to Th0 cells (Figure 5C&5D). Importantly, the same result was observed with BDC2.5 TCR transgenic T cells, confirming that this observation was not restricted to insulin specificity or retrogenic TCR expression (Figure 5E&5F). To investigate whether increased tetramer binding can be translated into a functional advantage for Tregs, we stimulated TCR transduced aTregs and Tem cells with plate-bound Ins-tet. Tregs started with a higher background level of Nur77-GFP and showed accelerated kinetics of response to antigen compared to Tem (Figure 5G&5H). Overall, our data suggest that in addition to the TCR-intrinsic parameters, such as affinity, Tregs possess cell-intrinsic mechanisms to boost their antigen recognition capacity. This observation provides a possible explanation for why we did not observed stronger insulin reactivity from TCRs expressed by islet-infiltrating Tregs sorted on tetramer, even though islet Tregs show higher frequency of tetramer+ cells. The ability of Tregs to increase their sensitivity for antigen could also allow their recruitment into the tissue and provide them with a competitive advantage against effector T cells expressing similar affinity TCRs.

### Low reactivity TCR-Tregs show reduced accumulation in islets, stability and impaired function upon adoptive transfer

To compare the function of Tregs expressing different TCRs, we tested Treg function against a single effector. Tregs were isolated from spleens and ndLNs of TCR Rg mice and transferred into NOD.TCRαKO female recipients together with Teffs that expressed a highly pathogenic insulin specific TCR (4-8) (Figure 6A)^28^. Teff TCR (TE-4), high reactivity Treg TCR (TR-2) as well as control TCR 4-8 expressing Tregs showed effective regulation of 4-8 Teffs and protection against diabetes. However, the low reactivity Treg TCR (TR-13) showed no protection (Figure 6B). GFP and Ametrine fluorescence markers were used to track donor Teffs and Tregs, respectively. Significantly lower numbers of Foxp3+Ametrine+ cells were recovered from spleens and islets of TR-13 Treg recipients (Figure 6C-6E), suggesting that TR-13 Tregs had impaired survival capacity and were unable to accumulate in pancreatic islets. Meanwhile, in TR-13 Treg recipients, few islet-infiltrating Ametrine+ cells maintained Foxp3 expression, suggesting that TR-13 Tregs were also unstable (Figure 6F). Interestingly, when TE-4, TR-2, and TR-13 Foxp3-Teff cells were transferred into a separate group of recipient mice, comparable numbers of donor Teffs were recovered from spleen and ndLNs across all TCRs (Figure S8A). The impaired survival observed in TR-13 with low insulin reactivity appeared to be Treg specific, suggesting that Tregs have more stringent TCR requirements for survival and function.

**Figure 6.**
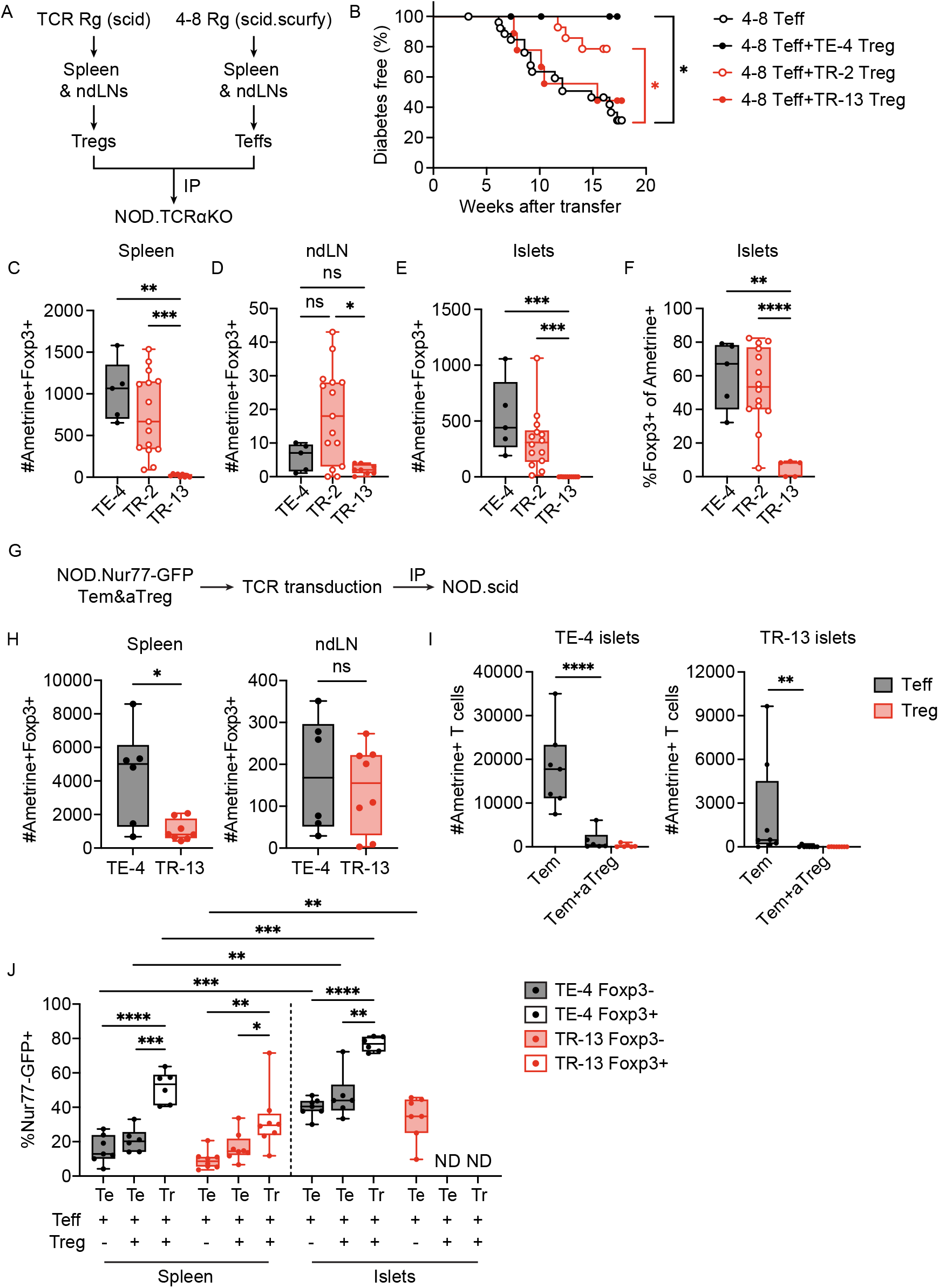
Treg TCR with low insulin reactivity showed reduced stability, accumulation in pancreatic islets and protective function. **(A)** CD4+ Tregs (Ametrine+GITR^hi^CD25+) were sorted from spleens and ndLNs of TCR Rg mice generated using NOD.scid BM. 4-8 TCR Rg mice were generated using NOD.scid.scurfy BM, and CD4+ Teffs cells (TCR-vector GFP+) were sorted from spleens of 4-8 Rg mice and transferred (IP) alone or with Ametrine+Tregs into NOD.TCRαKO recipients (Teff:Treg=5:1). Recipients were monitored for diabetes for up to 16 weeks. **(B)** Diabetes incidence: 4-8 Teff alone (n=27); 4-8 Teff+TE-4 Treg (n=8); 4-8 Teff+TR-2 Treg (n=14); 4-8 Teff+TR-13 Treg (n=9). *, p ≤ 0.05; ns > 0.05 by log-rank test. **(C-E)** Numbers of donor Tregs in spleens, ndLNs and pancreatic islets of recipient mice. **(F)** Percentages of Foxp3+ cells within Ametrine+ donor CD4+ T cells recovered from pancreatic islets. **(G)** aTreg and Tem cells were sorted from spleens and ndLNs of NOD.Nur77-GFP mice and transduced with indicated TCRs. Transduced (Ametrine+) cells were sorted and transferred (IP) into NOD.scid female recipients. Recipients were sacrificed for analysis two weeks after transfer. Data were pooled from two independent experiments. **(H)** Numbers of Ametrine+ Tregs recovered from spleen and ndLNs. **(I)** Numbers of Ametrine+ donor T cells in pancreatic islets. **(J)** Nur77-GFP expression in spleen and islets. Complete statistical alnalysis result is shown in Supplemenytary Fiigure 7B ****, p ≤ 0.0001; ***, p ≤ 0.001; **, p ≤ 0.01; *, p ≤ 0.05; ns > 0.05 by (C-E, I) Kruskal-Wallis test followed by Dunn’s test, (F, J) Welch ANOVA followed by Benjamini, Krieger and Yekutieli’s test and (H) Mann-Whitney test. Populations with less 10 cells and outliers were excluded from analysis using ROUT method (Q=1). See also Figure S8.

To address whether our *in vivo* observations were directly imposed by TCR-intrinsic parameters or shaped by TCR-specific development and peripheral stimulation in single TCR mice, we TCR-transduced primary polyclonal aTregs and Tem cells isolated from spleens and ndLNs of NOD.Nur77-GFP mice and co-transferred cells into NOD.*scid* female recipients (Figure 6G). Consistent with TCR Rg Treg transfers, significantly lower numbers of TR-13 Tregs were recovered from spleens but not ndLNs of recipient mice (Figure 6H), indicating that insulin reactive Treg survival or expansion is continuously regulated by the level of TCR activation. To address whether low reactivity Treg TCRs were sufficient to impart suppressive capacity onto fully mature and functional polyclonal Tregs, we tested the ability of TCR transduced Tregs to suppress Teffs *in vivo*. Both high and low reactivity TCR transduced aTregs inhibited islet-infiltration of Tem cells that expressed the same TCR (Figure 6I). These observations show that Treg functional capacity varies depending on the affinity of the responder T cells. In other words, low affinity Tregs could suppress low affinity effectors, but are insufficient against highly potent effector T cells. This might explain why Tregs ultimately fail against high affinity neo-antigen reactive T cells, such as hybrid insulin peptides, since neo-antigen reactivity by definition is not curbed during thymic negative selection. An increase in Nur77-GFP expression from spleen to islets was observed in TE-4 Tem cells, TE-4 aTregs and TR-13 Tem cells (Figure 6J, Figure S8B), validating antigen-specific induction of TCR signaling. Significantly higher level of Nur77-GFP expression was observed in Tregs compared with Teffs that expressed the same TCR (Figure 6J), which mirrors the increased tetramer staining and Nur77-GFP expression we observed in retrogenic and TCR transduced polyclonal Tregs (Figure 5). In combination, elevated Treg TCR signal strength *in vivo* and Treg increased capacity to bind pMHC ligands, as determined by tetramer staining, suggest that Tregs can be more effective at responding to antigen *in vivo* to effectively suppress Teffs with similar TCR specificity and affinity.

## Discussion

The two-step instructive model suggests that a strong, transient TCR signal is favored for Treg development^44,45^. This model is supported by the finding that Tregs express TCRs with stronger self-reactivity than Teff cells.

Ablation of cognate antigen expression in the thymus inhibited Treg, but not conventional T cell development, suggesting the difference in TCR repertoire between the two populations to be primarily driven by agonist selection of Tregs^46^. Meanwhile, in a fixed pMHC antigen model ubiquitous expression of antigen selected Tregs and Teffs with similar TCRs, suggesting that self-reactive Tregs and Teffs could traverse thymic selection with similar TCR affinities^10^. Interestingly, the difference in TCR repertoire in the ubiquitous antigen model was more pronounced at the higher end of the TCR affinity spectrum, implying that negative selection of Teffs drove the difference in repertoires^10^. Consistently, unlike tissue restricted antigen modelled by Cre recombinase driven by insulin promoter, ubiquitously expressed Cre antigen was shown to drive deletion of conventional T cells instead of Treg generation^47^. Analogous observations were obtained in a fixed-TCR system, where ectopic over-expression of insulin increased negative selection of high affinity, but not low affinity T cells, while both TCRs supported Treg development^29^. Our initial observations showing increased expression of CD5 and Nur77-GFP on Tregs in wild type NOD mice and in TCR limited P2-TCRα mice were consistent with the two-step instructive model (Figure 1C&1D, Figure 3B&3C). We also observed a decrease in Treg development in mice with mutated target insulin epitope, indicating a critical role for specific self-antigen in Treg development (Figure S4C)^6^. However, the shorter CDR3β expressed by islet insulin specific Teffs, which is favored during thymic selection^48,49^, was also not observed in repertoires generated in wild-type insulin knockout mice (Figure 3G, Figure S4E). We also did not observe a stronger reactivity to islet autoantigens in Treg TCRs (Figure S1, Figure 3I). Together, our data suggest that positive selection of Tregs and deletion of conventional T cells simultaneously shape the divergent Treg and Teff TCR repertoires. Since both Tregs and Teffs can be selected with relatively high affinity TCRs, but prefer distinct TCR characteristics, it is possible that divergence in TCR repertoire is driven by different niches for Treg and Teff development, potentially instructed by secondary signals provided by distinctive APCs.

Despite most of the islet-infiltrating CD4+ T cells in P2-TCRα Rg mice showing insulin reactivity in the micropipette adhesion analysis, many of the isolated TCRs failed to elicit significant response when re-expressed and tested for functional reactivity (Figure 3D&3I). However, TCR affinity does not always predict T cell functional outcome^50^. It is possible that for some antigens difference in Treg/Teff TCR repertoires might underlie structural differences of how Treg TCRs engage pMHC ligands^51^. Biophysical parameters and co-receptors can also influence signal transmission in T cells and T cell recruitment into immune response^52^. Our observations regarding higher level of TCR signaling and tetramer-binding in Tregs expressing the same TCR as effector T cells is intriguing and suggests cell-intrinsic capacity of Tregs to augment responses to TCR/pMHC interaction (Figure 5). The cell-intrinsic mechanisms that support higher Treg sensitivity for pMHC ligands is unclear, but tetramer staining potentially suggests differences in cell-membrane organization of TCR complexes on Tregs. How this Treg quality is translated into higher TCR signaling in response to self-ligand *in vivo* is unknown, but it is tempting to postulate that it supports an increased ability of Tregs to compete for limiting amounts of pMHC ligands. In support of this, we observed that *in vitro* Tregs responded to pMHC antigens with faster kinetics (Figure 5H). However, while low affinity TR-13 Tregs were able to infiltrate pancreatic islets (Figure 4C) and to compete for ligand as indicated by higher Nur77-GFP compared to TR-13 Teffs (Figure 6J), TR-13 Tregs failed to suppress higher affinity 4-8 TCR effectors (Figure 6B). Based on our observations, we propose that ubiquitous expression of insulin allows enlargement of the thymic Treg niche to support development of relatively lower affinity Tregs. Following selection, Treg-intrinsic capacity to compete for pMHC ligand permits recruitment of low affinity Tregs into the autoimmune response. Since low affinity Tregs, such as TR-13, are not as robust at suppressing high affinity Teffs, the balance of Treg/Teff TCR affinities may be shifted towards a loss of a Treg advantage in the context of ubiquitous self-antigens.

Tregs were previously reported to have dampened TCR signaling, as measured by maximum Erk phosphorylation, compared to naïve T cells^53^. We did not observe a significant difference in maximum Nur77-GFP level between aTregs and Tems (Figure 5H). It is possible that difference in the magnitude of upstream signaling pathway activation is not directly translated to downstream gene expression. Our observations showing increased basal and *in vivo* induced Nur77-GFP expression in Tregs are consistent with a recent study that demonstrated a Treg specific pattern of TCR signaling component activation, including a higher level of ITAM phosphorylation at a steady state^54^. Further investigation is needed to resolve the relationship between TCR signaling regulation, Treg sensitivity to antigen and how downstream signaling ultimately influences Treg suppressive function.

It is unclear whether the initial TCR signal during thymic selection that drives the Treg development is initiated by cell-intrinsic increased capacity for TCR signaling and whether it is directly driven by Foxp3. Our observations that iTregs bind pMHC tetramers at a higher level compared to *in vitro* generated Th0 cells suggest that expression of Foxp3 can impact T cell sensitivity for antigen, irrespective of the quality of the endogenous TCR and signals acquired during thymic selection (Figure 5C-5F). On the other hand, CD25+Foxp3-thymic Treg precursors already exhibit high levels of TCR signaling prior to upregulation of Foxp3^55^. CD25-Foxp3+ and CD25+Foxp3-thymic Treg precursors exhibit distinct TCR repertoires, suggesting development in response to different peptide ligands^55^. Expansion of thymic antigenic niche preferentially leads to CD25-Foxp3+ Treg pathway of development^31^, suggesting that low affinity self-reactive TCRs associated with increased antigenic niche follow the CD25-Foxp3+ trajectory. Therefore, Foxp3 expression might be an important factor for increased Treg sensitivity for antigen.

In conclusion, our study provides a detailed analysis of TCR repertoires and TCR characteristics exhibited by Tregs recruited into the spontaneous autoimmune response in the context of a susceptible MHC allele. While some observations were consistent with previous studies, such as higher expression of CD5 and Nur77-GFP in Tregs, this increase in TCR signaling was not reflected in Treg TCR reactivity for insulin. Moreover, insulin specific Treg TCRs failed to support better Treg development or function in inhibiting diabetes compared to Teff TCRs. This surprising diminished suppressive potency of Treg TCRs may contribute to the pathogenesis of diabetes in NOD mice, and suggests that Treg TCRs may not be the preferred choices to be used in TCR-redirected Treg therapy in human T1D. Overall, our study puts an emphasis on the relative capacity of Treg and Teff cells to respond to self-antigen as the deciding factor in tolerance vs autoimmunity.

## Materials and Methods

### Mice

NOD/ShiLtJ (NOD), NOD/ShiLt-Tg(Foxp3-EGFP/cre)1cJbs/J (NOD.Foxp3-GFP), NOD.Cg-*Prkdc*^*scid*^/J (NOD.scid) and NOD.*Tcra*^*tm1Mjo/DoiJ*^ (NOD.TCRαKO) mice were purchased from Jackson Laboratory. B6.Nur77^GFP^ mice ^27^ were backcrossed to NOD at least 10 generations, and subsequently crossed to NOD.*scid* (^28^. NOD.TCRαKO.Ins1KO.Ins2KO.InsY16A mice were generated by crossing NOD.Cg-Tg(Ins2*Y16A)3Ell *Ins1*^*tm1Jja*^ *Ins2*^*tm1Jja*^/GseJ mice ^34^ with NOD.TCRαKO mice ^6^. NOD.TCRαKO.Foxp3-GFP mice were generated by crossing NOD.Foxp3-GFP mice with NOD.TCRαKO mice. All mice were housed under specific-pathogen-free conditions in Baylor College of Medicine and University of Utah facilities. All experiments were performed in accordance with Institutional Animal Care and Use Committee protocols at Baylor College of Medicine and University of Utah.

### Flow cytometry and FACS

Surface staining of cell lines and primary cells was performed in PBS with 3% FBS and 0.05% sodium azide. Fc receptor was blocked by incubating with TruStain FcX reagent (BioLegend, 101320). Viable cells were identified using Zombie Dyes (Biolegend, 423109&423107). For intracellular staining, cells were fixed using the eBioscience™ Foxp3 / Transcription Factor Fixation/Permeabilization (00-5521-00) or 2% methanol free paraformaldehyde (Thermo Scientific, 28908) at room temperature for 30min. Fixed cells were washed twice before intracellular staining in eBioscience™ Permeabilization Buffer (00-8333-56) on ice overnight. Flow cytometry analyses were performed on LSR Fortessa II (BD Biosciences) or Aurora (Cytek). FACS was performed using BD FACSAria II/III or Sony MA900 cell sorter. Data were analyzed with FlowJo software (TreeStar). Monoclonal antibodies against following markers were used: CD4 (GK1.5), CD8α (53-6.7), CD5 (53-7.3), CD3 (145-2C11), TCRβ (H57-597), IFNγ (XMG1.2), CD62L (MEL-14), CD44 (IM7), CD25 (PC61), GITR (TYGITR 765), CD45.1 (A45), CD45.2 (104), CD11b (M1/70), CD11c (N418), B220 (RA3-6B2), PD-1 (29F.1A12), CD73 (TY/11.8) from Biolegend, Foxp3 (FJK-16s), CD11b (M1/70), B220 (RA3-6B2), γδTCR (UC7-13D5), TER-119 (TER-119), CD49b (DX5) from eBioscience, and TCR Vβ4 (REA729) from Miltenyi Biotec. InsB:9-23 E21G/R22E, InsB:9-23 R22E and 2.5HIP tetramers were obtained from the National Institutes of Health (NIH) Tetramer Core Facility. Unless otherwise stated, tetramer staining was performed by 1hr on ice incubation.

### Peptide

Insulin B:9-23 wild-type (SHLVEALYLVAGERG), R22E (SHLVEALYLVAGEEG) and E21G/R22E peptides (SHLEVALYLVAGGEG) were purchased from GenScript. 2.5HIP (DDLQTLALWSRMDQLDD) was purchased from Genemed Synthesis Inc.

### Generation of TCR retrogenic mice

TCR retrogenic mice were generated as previously described ^19^. Briefly, 5-fluorouracil (NDC, 63323-117-51) was given intraperitoneally to bone marrow donor mice (0.15mg/g body weight). Donor mice were sacrificed 3 days later and isolated BM cells were cultured in DMEM with 20% FBS and supplements, including non-essential amino acids (Quality Biological, 116-078-721), glutamine (Corning, 25-005-CI), sodium pyruvate (Corning, 25-000-CI), HEPES (Corning, 25-060-CI), beta-mercaptoethanol (Gibco, 21-985-023) and penicillin-streptomycin) (Corning, 30-002-CI), and cytokines (mSCF, IL-6, hIL-3). Retroviral supernatant was generated using TCR-vector transduced GP+E86 cells. BM cells were retrovirally transduced with retroviral supernatant twice with by spinoculation (2,500rpm, 1hr, 37degree). Transduced BM cells (2-4 million) were transferred intravenously into sub-lethally irradiated (500 rads) NOD.TCRαKO mice. Unless otherwise stated, NOD.*scid* mice were used as bone marrow donors.

### Assessment of diabetes

TCR retrogenic mice were monitored for diabetes weekly, starting from 5wk after BM transfer. Diabetes onset was defined as blood glucose ≥ 400mg/dl or ≥ 300mg/dl for two consecutive days.

### Pancreatic Islets isolation

T cells that infiltrate pancreatic islets were isolated after intra-bile duct injection and digestion with collagenase IV (Worthington Biochemical) ^28^. Perfused pancreata were incubated at 37°C for 30min and washed twice with 5%FBS+HBSS (Corning). Islets were hand-picked and dissociated at 37°C in 1ml of cell dissociation buffer (Gibco) for 15min. Dissociated islets were washed with HBSS before proceeding to analysis.

### TCR repertoire sequencing

For single-cell RNAseq and TCRαβ repertoire analysis, pancreatic islets were harvested from four 12-week-old female NOD mice and labelled individually with anti-TCRβ (BioLegend, allophycocyanin, H57-597, 1:300), anti-CD4 (BioLegend, TotalSeq-C0001, RM4-5, barcode-AACAAGACCCTTGAG, 1:1000), anti-CD8b-biotin (BioLegend, YTS156.7.7, 1:8000) primary antibodies followed by secondary labelling with PE SA(BioLegend, TotalSeq-C0951, barcode-AACCTTTGCCACTGC, 1:8000). Dead cells were excluded during sorting using DAPI staining (Thermo Fisher, D1306). TCRβ+ cells were sorted from the islets on an Aria sorter and resuspended at 1000 cells/μl in PBS with 0.04% BSA.

Libraries were generated using Chromium Next GEM Single Cell 5’ Reagent Kits v2 (Dual Index) with Feature Barcode technology for Cell Surface Protein and Immune Receptor Mapping. Cells were then loaded into the 10X Controller. The resulting GEMs were transferred to new tubes for GEM-RT. After the GEM-RT incubation the samples were cleaned-up and readied for cDNA amplification. Amplified cDNA was then size-selected using SPRIselect Beads (BECKMAN COULTER). The resulting cDNA was used to generate the various library types; V(D)J TCR, 5’ Gene Expression (GEX) and Cell Surface Receptor. Sequencing of Libraries was completed at the Genomics and Microarray Core, University of Colorado Anschutz Medical Campus, using a full lane on an Illumina NovaSeq 6000 using standard protocols for 10X requirements. scRNAseq and CITE-seq data were analyzed with Seurat as follows. scRepertoire was used for scTCRseq analysis of CD4 Teff and Treg subsets.

For antigen agnostic T cell isolation and TCR cloning (Table I) from pancreatic islets, single cells were sorted from islets of one 13 week pre-diabetic NOD.Foxp3-GFP mouse based on GFP expression. Sequences of Treg (CD4+CD3+GFP+) and Teff (CD4+CD3+GFP-) TCRs were obtained using a previously described multiplex-nested PCR based method ^56^. Briefly, reverse transcription was performed on sorted single cells using reverse primers for TCR alpha and beta chain constant regions, which was followed by a round of PCR using a mix of 44 and 40 TCRα and TCRβ variable region primers respectively, and the corresponding constant region external primers. PCR products were subcloned into retroviral vector, sequenced, and analyzed with reference to the IMGT database ^57^.

For TCRβ library preparation for ImmunoSEQ, CD4+ Teffs (Foxp3-GFP negative) and Tregs (Foxp3-GFP positive) isolated from pancreatic islets of P2-TCRα retrogenic mice were stained with PE and allophycocyanin conjugated insulin B:9-23(E21G/R22E) tetramer. Tetramer positive cells were sorted into ATL buffer (Qiagen) and stored at -80°C. Splenocytes of P2-TCRα retrogenic mice were enriched for CD4+ cells using CD4 microbeads (Miltenyi Biotec, 130-117-043). Teffs (≤ 140,000 cells per mouse) and Tregs were sorted from enriched cells into ATL buffer and stored at -80°C without tetramer staining. Genomic DNA was purified from frozen samples using QIAamp DNA Micro Kit (Qiagen). Sequencing data were analyzed using the ImmunoSEQ analyzer (version 3.0) and vegen and vvenn R packages.

### Functional avidity measurements

TCR functional avidity was measured using TCR transduced 4G4.CD4+ thymoma cells ^28^ or *in vitro* expanded retrogenic CD4+ T cells. TCR negative 4G4.CD4+ cells were transduced with TCR expression vectors by two consecutive spinoculations (1800rpm, 20°C, 90min) performed one day apart. Transduced cells were purified by CD3ε+ MACS enrichment. A total of 50,000 TCR+4G4.CD4+ cells were stimulated with plate-bound tetramer or peptide and 25,000 I-A^g7^ expressing M12.C3 lymphoma cells. After 24hr, IL-2 concentration in supernatant was measured by ELISA. For IL-2 ELISA, 96-well high adhesion plates (VWR, 490012-252) were coated with IL-2 antibody (Biolegend, 503702) and blocked using 1% BSA (Fisher Scientific, BP9703-100). Biotinylated IL-2 antibody (eBioscience, 13-7021-85), SA-HRP (Biolegend, 405306) and TMB (BD Biosciences, 555214) were used for detection. Reaction was stopped by adding sulfuric acid and absorbance at 450nm was then measured. For TCR functional avidity measurement using retrogenic T cells, spleens of TCR Rg mice were dissected, homogenized and CD4+ T cells were isolated using MACS enrichment. Purified CD4+ T cells were expanded by stimulating T cells with phorbol myristate acetate (PMA, Sigma-Aldrich, 10ng/ml) and ionomycin (Sigma-Aldrich, I0634, 1*μ*g/ml) for 48hr and followed by expansion with 1000U/ml hIL-2 (PeproTech) for two weeks. Expanded T cells were stimulated with peptide and DCs obtained from B16.Flt3l melanoma immunized NOD.TCRαKO mice ^30^. Cells were stimulated for 5hrs in the presence of brefeldin A and monensin (Biolegend). Intracellular IFNγ was measured by flow cytometry.

### 2D affinity measurement

TCR two-dimensional affinity was measured using the micropipette adhesion frequency assay which was previously described in detail ^23,58^. Briefly, biotin-tagged peptide-MHC monomers were coated on biotinylated human red blood cells (RBCs) via biotin-streptavidin chemistry. Individual coated RBCs were used as antigen-presenting cells and were mechanically controlled to approach, contact (for 2s) and retract from CD4+ T cells placed on opposing micropipettes. This cycle was performed 50 times per T cell/RBC pair. The presence of adhesion (indicating TCR–pMHC ligation) was readout as elongation of the RBC membrane during retraction and adhesion frequency (Pa) was calculated as the number of adhesion events divided by total cycle number.

Cells with Pa greater than the cutoff for nonspecific binding (to CLIP:I-A^g7^, nominally 10%) were deemed as antigen-specific. For each cell pair, 2D affinity (A_c_K_a_) was calculated using the following equation:

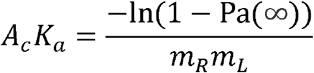

Pa(∞), adhesion frequency at an equilibrium contact time (≥2 s).

m_R_ and m_L_, desities of the receptor (TCR) and ligand (pMHC), respectively.

### Treg and Teff cell purification, T cell TCR transduction and *in vitro* T cell differentiation

CD4+ T cells were isolated from spleens and non-pancreatic draining lymph nodes (ndLN) by MACS enrichment (Miltenyi Biotec). Treg and Teff cells were then sorted from CD4+ T cells (Teff: CD4+CD5+CD25-GITR^lo^; Treg: CD4+CD5+CD25+GITR^hi^). Post-sort purity was determined by intranuclear Foxp3 staining (≥95% for polyclonal primary T cells and retrogenic Teffs; ≥ 90% for retrogenic Tregs). For TCR transduction, sorted cells were stimulated with anti-CD3 and anti-CD28 coated dynabeads (Thermo Fisher, beads:cells=1:1), and were transduced retrovirally by spinoculation (2,500 rpm, 37°C, 1hr) at 24hr and 48hr after stimulation. Teffs were supplemented with 50U/ml hIL-2 (PeproTech) and Tregs with 1000U/ml hIL-2. For *in vitro* differentiation, naïve CD4+ T cells (CD25-CD44-CD62L+) were sorted, stimulated with plate-bound anti-CD3 (Biolegend, 1μg/ml) and anti-CD28 (Biolegend, 2μg/ml) for 48hr, and were cultured with 50U/ml hIL-2 (Th0) or 100U/ml hIL-2 and 5ng/ml murine TGFβ (R&D Systems, 7666-MB/CF) for 4-5 days.

### Statistical analysis

All data analysis was performed using GraphPad Prism 10 (GraphPad Software). Statistical significance between sample means were indicated as such: *P < 0.05; **P < 0.01; ***P < 0.001; ****, p ≤ 0.0001.

## Data and materials availability

The datasets generated during and/or analyzed during the current study are available from the corresponding author on reasonable request.

## Acknowledgements

The authors thank the NIH Tetramer Core Facility (Atlanta, GA) for providing the 2.5HIP:I-A^g7^, insulin B:9-23(E21G/R22E):I-A^g7^ and insulin B:9-23(R22E):I-A^g7^ tetramers. The authors also thank the Cytometry and Cell Sorting Core at Baylor College of Medicine (Houston, TX) and the Flow Cytometry Core at University of Utah (Salt Lake City, UT) for technical support. This work was supported by the NIH (AI125301 to M.Be. and DK114456 to M.L.B.), and The Robert and Janice McNair Foundation.

## Author contributions

Conceptualization: YJ, MLB, MB. Methodology: YJ, YK, DA, BL, EK, MS, MLB, MB. Investigation: YJ, YK, DA, BL, MB. Visualization: YJ, DA, MB. Funding acquisition: MLB, MB. Project administration: YJ, MB. Supervision: BE, MLB, MB. Writing – original draft: YJ, DA, MB. Writing – review & editing: YJ, DA, BL, BE, MLB, MB.

