## Supplementary Figures and tables for "Increased TCR signaling in regulatory T cells is disengaged from TCR affinity"

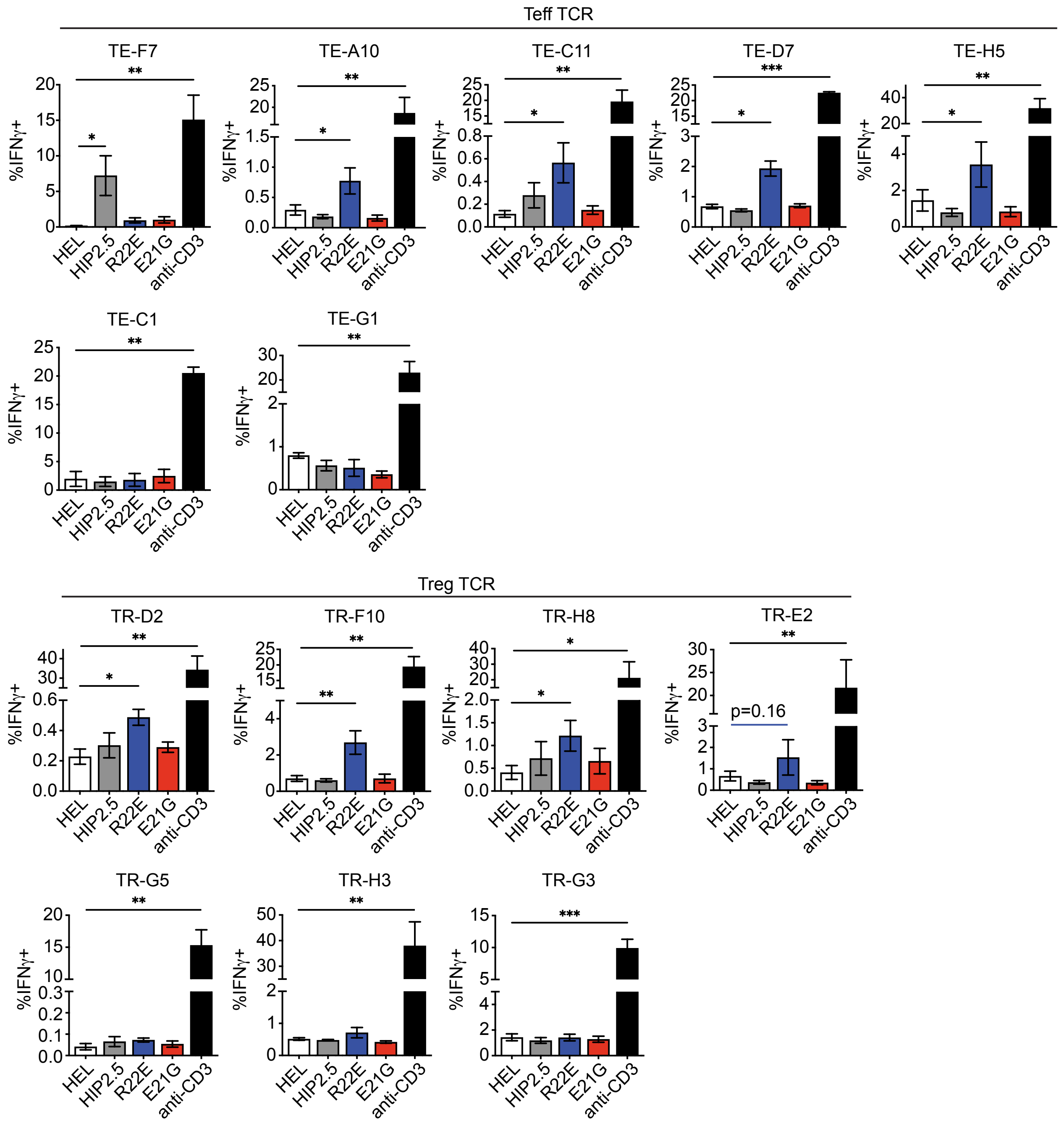


**Supplementary Figure 1. Pancreatic islet infiltrating Tregs do not express TCRs with higher affinity compared with Teffs.** CD4+ T cells were isolated from spleens of single TCR Rg mice, stimulated with PMA and ionomycin, and expanded *ex vivo* with 1000U/ml hIL-2 for 2 weeks. Expanded CD4+ T cell clones were then stimulated with DCs and indicated peptides (50µM) for 5hr in the presence of BFA and monensin. Intracellular IFN was measured as readout of T cell activation. Data are plotted as means ± SEM and are pooled from at least 3 independent experiments. HEL, hen egg lysozyme, epitope 11-25. ***, $p \leq 0.001$; **, $p \leq 0.01$; *, $p \leq 0.05$; ns > 0.05 by one-tail paired t test (n=3-7).

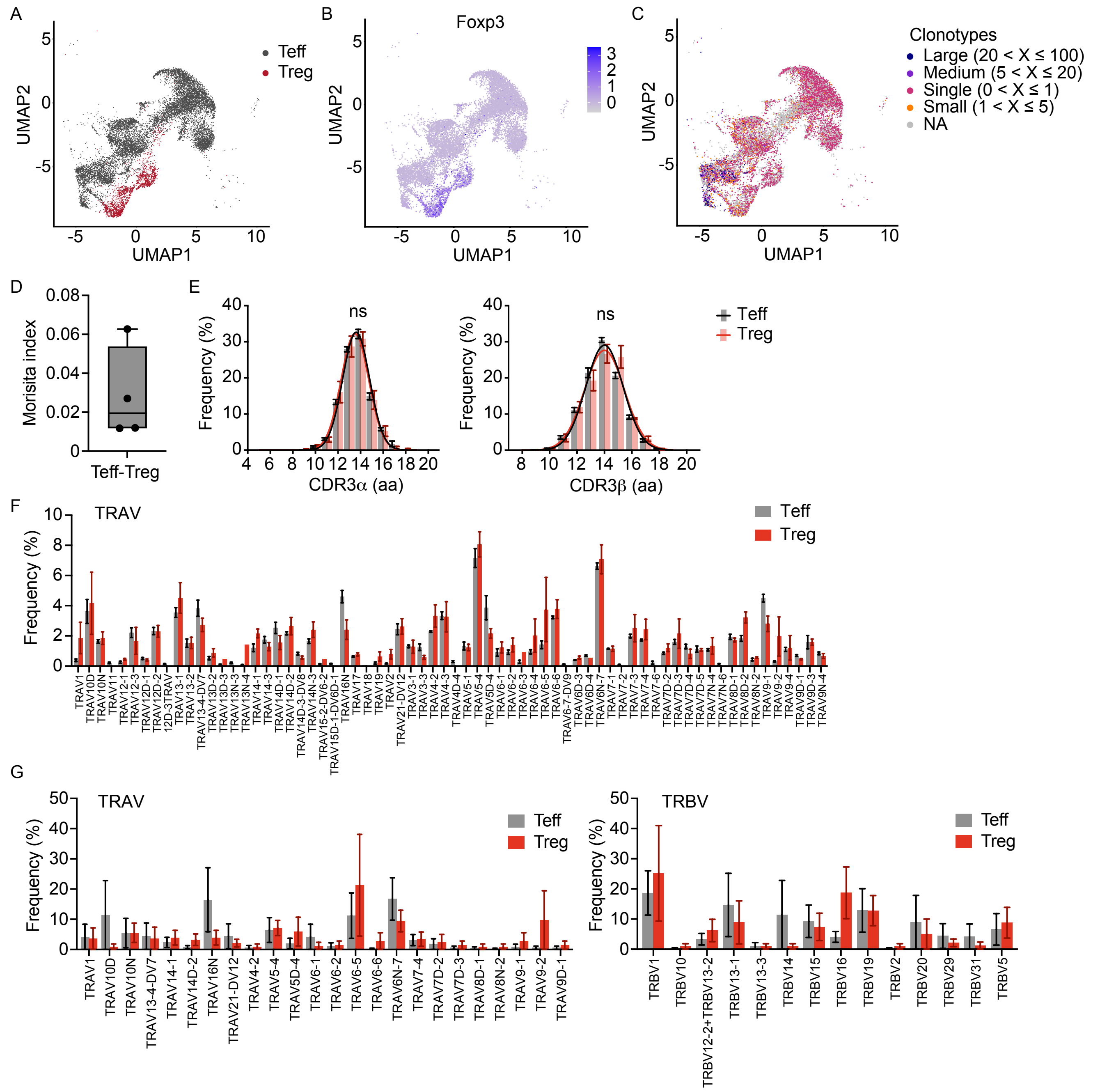


**Supplementary Figure 2. Single cell RNAseq analysis of TCRaβ repertoires of Teffs and Tregs isolated from pancreatic islets of four pre-diabetic NOD females.** Single cell RNAseq analysis of TCRaβ repertoires of Teffs and Tregs isolated from pancreatic islets of four 12-week-old pre-diabetic NOD females. **(A, B)** Tregs and Teffs were distinguished using UMAP based analysis and corresponding Foxp3 expression. **(C)** TCR clonotype (unique CDR3 amino acids sequences) distribution. **(D)** Similarity between Treg and Teff TCR repertoires of the same mouse was measured by Morisita index. **(E)** Distributions of CDR3α and CDR3β length. aa, amino acids. **(F)** TRAV gene usage. **(G)** V segment gene usage of TCRs identified from both Treg and Teff repertoires. Data are plotted as means ± SEM (n=4) and analyzed by (E) extra sum-of-squares F test and (F&G) multiple paired t test. ns > 0.05.

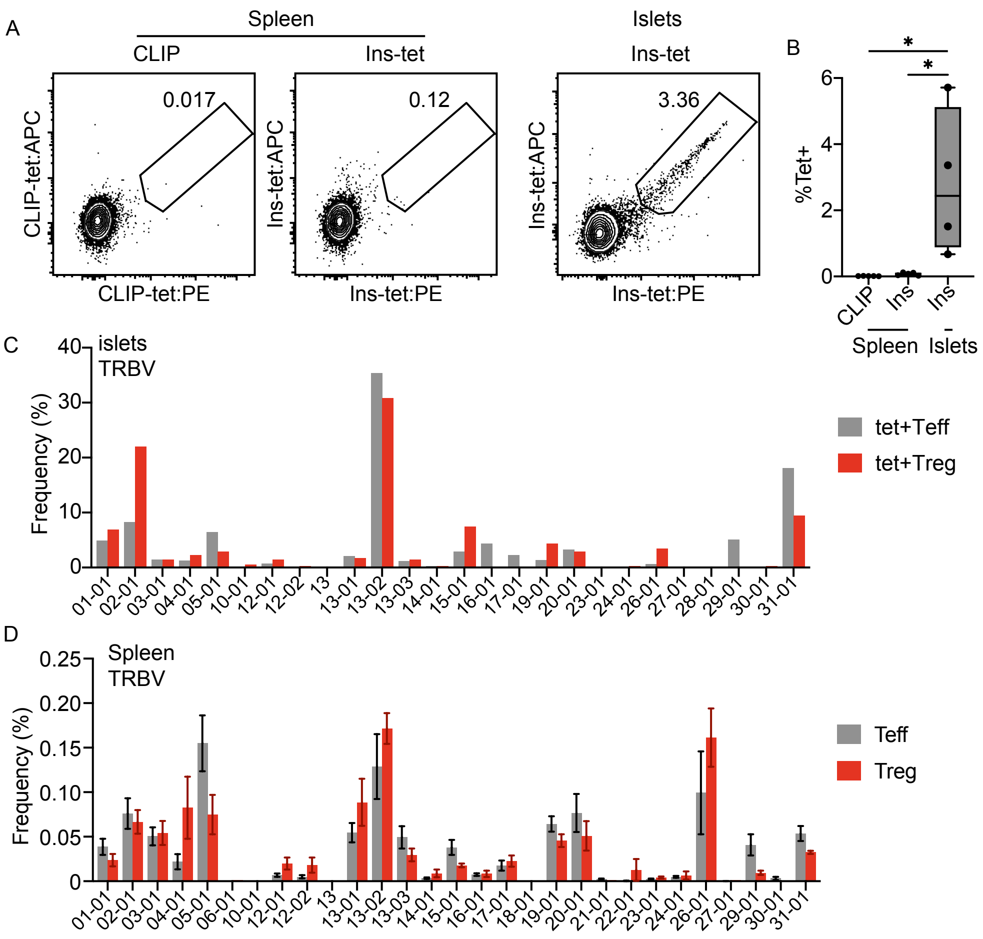


**Supplementary Figure 3. Insulin specific CD4+ T cells were enriched in pancreatic islets of P2-TCRa Rg mice. (A, B)** Splenocytes and pancreatic islets of P2-TCRα Rg females were stained with Ins-tet or CLIP:I-A$^{g7}$ tetramers. Frequencies of tetramer positive CD4+ T cells were shown. Gated on Zombie red-CD4+CD5+CD3+Ametrine+ cells. Spleen, n=5; Islets, n=4. **(C, D)** Ins-tet+ cells from pancreatic islets of 38 P2-TCRα mice were sorted, pooled and sequenced. Tregs and Teffs were also sorted from 5 individual spleens without tetramer staining and were sequenced separately. TRBV gene usage of (C) islet Ins-tet+ and (D) splenic repertoires. (B) One-way ANOVA followed by Holm-Sidak's test. (D) Data are plotted as means ± SEM (n=5) and analyzed by paired multiple t test. *, $p \leq 0.05$; ns > 0.05.

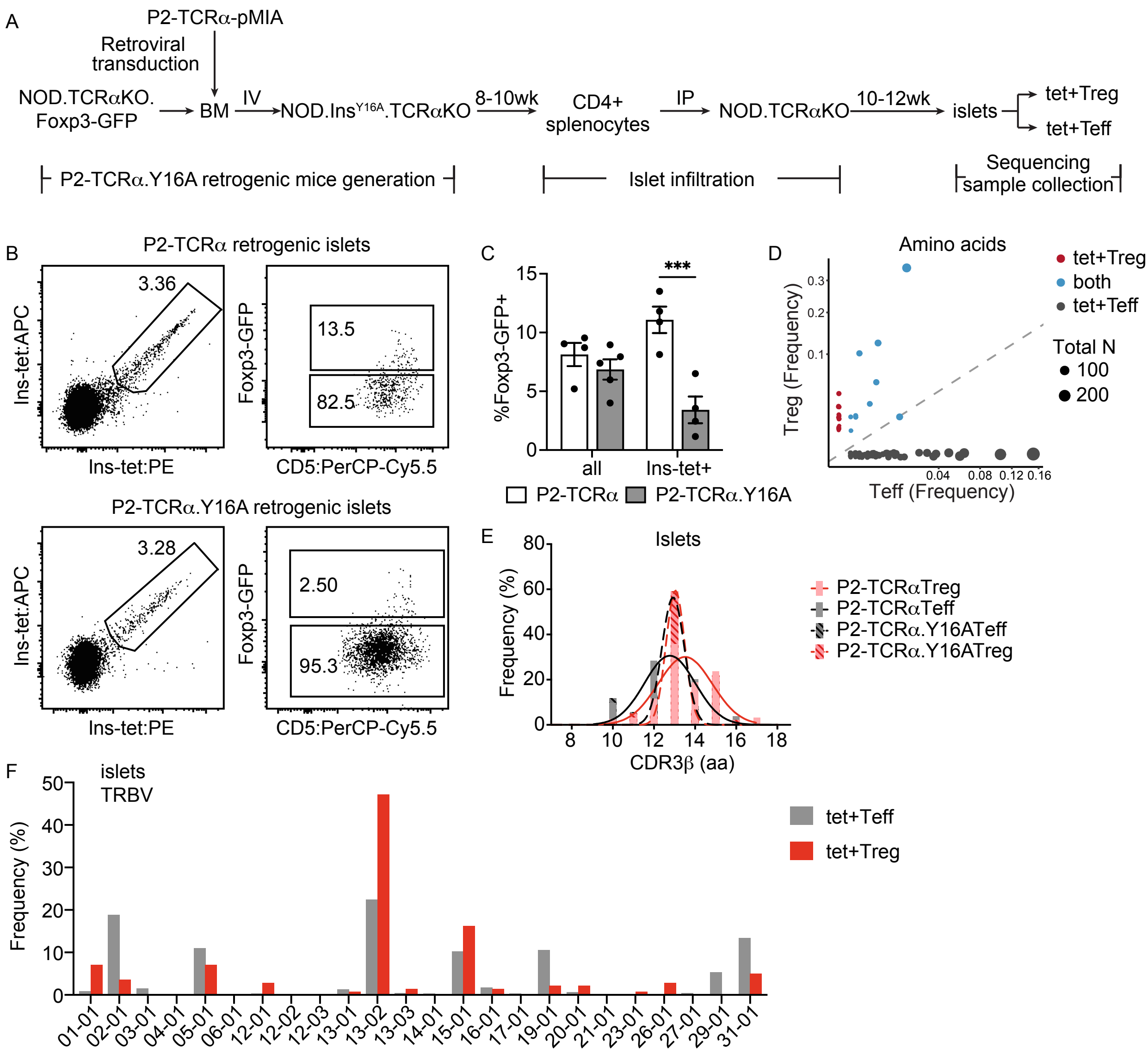


**Supplementary Figure 4. TCRβ repertoire analysis of islet Ins-tet+ Tregs and Teffs of P2-TCR α.Y16A mice. (A)** Diagram for P2-TCRα.Y16A retrogenic mice generation and Ins-tet+ TCRβ repertoire analysis. BM cells of NOD.TCRαKO.Foxp3-GFP mice were transduced with a retroviral vector that encodes the TCRα chain of TCR P2 and an Ametrine expression reporter. Transduced BM cells were transferred (IV) into NOD. $Ins^{Y16A}$.TCRαKO mice. Splenic CD4+ T cells were isolated from BM recipients 8-10wk after BM transfer and transferred (IP) into NOD.TCRαKO mice. 10-12 weeks later, pancreatic islet-infiltrating Treg and Teff cells were isolated and stained with Ins-tet. TCRβ repertoires of Ins-tet+ Tregs and Teffs were sequenced and analyzed. **(B, C)** Ins-tet staining and frequencies of Foxp3-GFP+ Tregs within Ins-tet+ populations or all islet CD4+ T cells (all). Data are plotted as means ± SEM (n=4 or 5) and analyzed by two-way ANOVA and Benjamini, Krieger and Yekutieli test. **(D)** Repertoire overlap between islet infiltrating Ins-tet+ Tregs and Teffs at amino acid level. Dot size represents size of clonotypes. **(E)** Distributions of CDR3β length. Extra sum-of-squares F test. **(F)** TRBV gene usage. ***, $p \leq 0.001$; ns > 0.05.

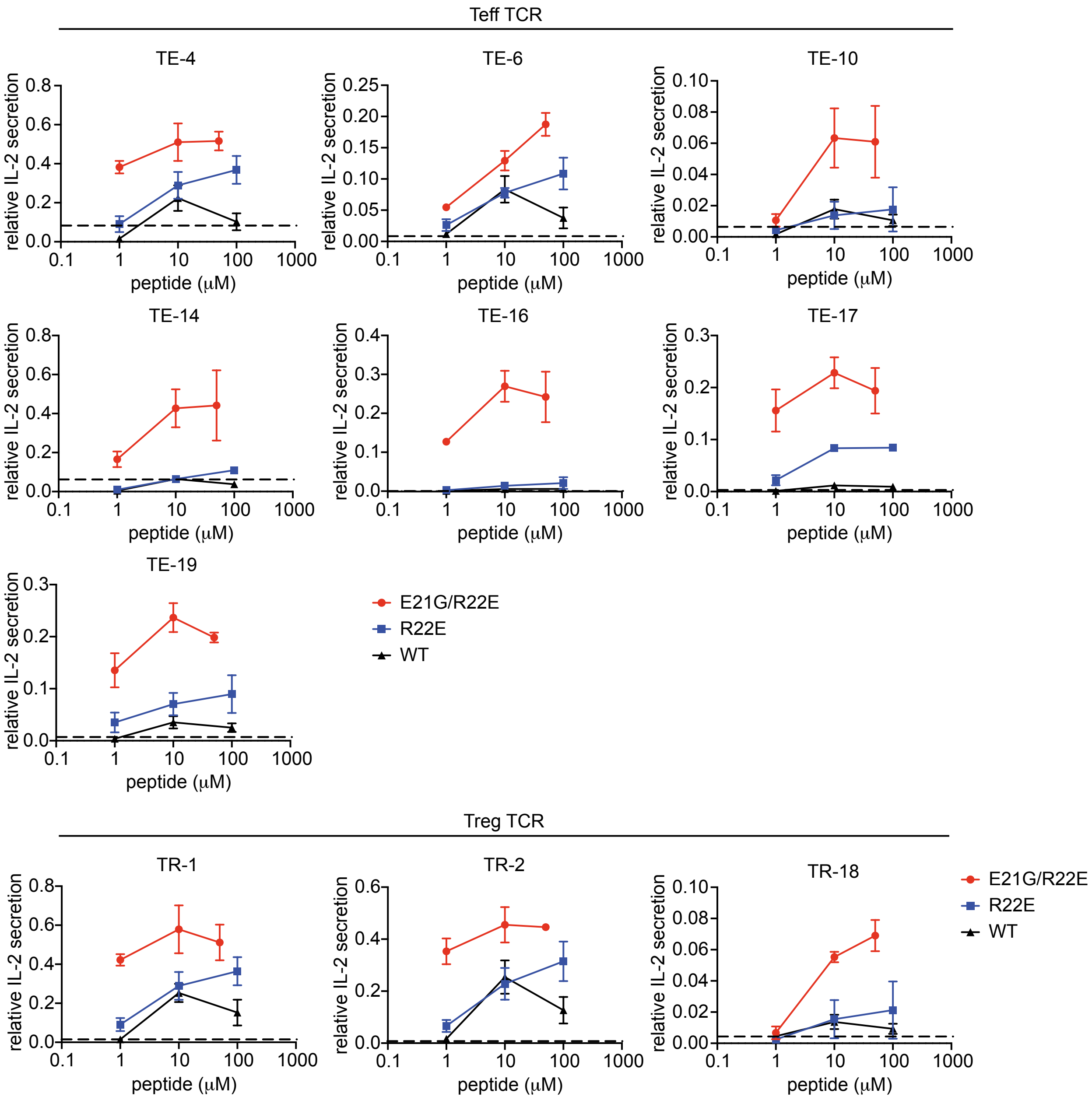


**Supplementary Figure 5. *In vitro* stimulation of 4G4 thymoma cells.** 4G4 thymoma cells expressing indicated TCRs were stimulated with insulin peptides and M12.C3.g7 lymphoma cells for 24hr. IL-2 concentrations in the supernatant were measured by ELISA and normalized to anti-CD3 positive control. Data are plotted as means ± SEM and are pooled from 3 independent experiments. Dash line, average of relative IL-2 secretion from unstimulated negative controls.

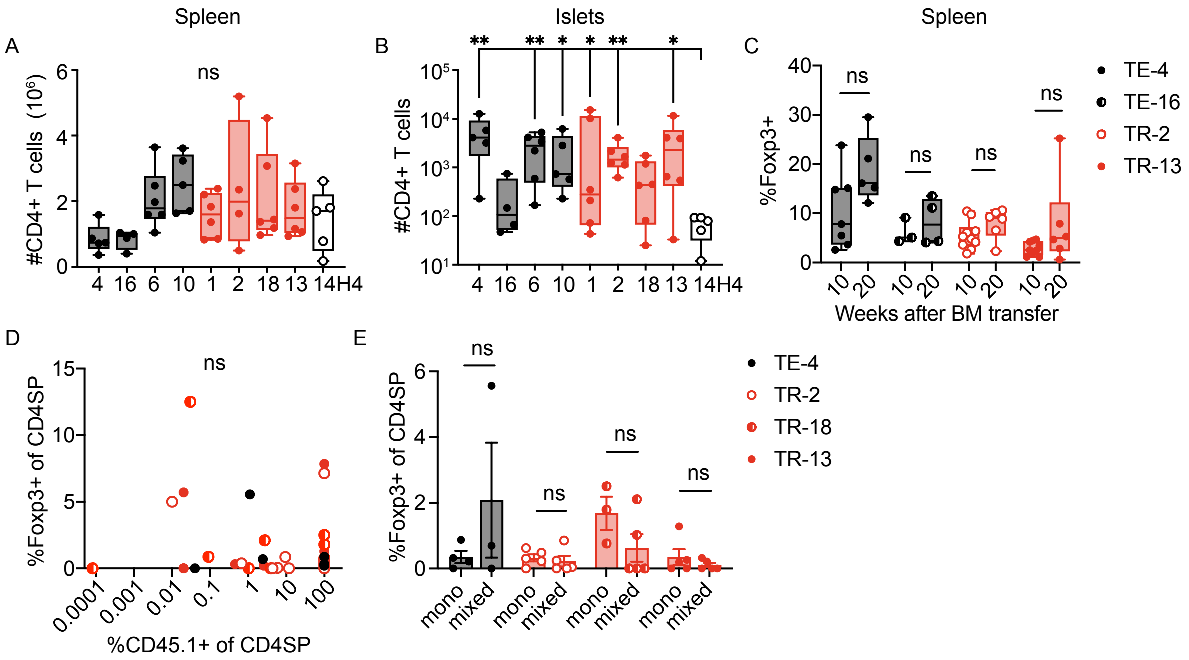


**Supplementary Figure 6**. **Supplementary data to Figure 4. (A-C)** Retrogenic mice expressing Teff (black), Treg (red) derived TCRs, or HEL specific 14H4 control TCR (white) were generated by transferring transduced NOD.*scid* BM (IV) into irradiated NOD.TCRαKO recipients. (A) Numbers of CD4+ T cells in spleens. (B) Numbers of Tregs in pancreatic islets. Mice were analyzed 20wk after BM transfer or at diabetes onset. (C) Frequencies of Tregs in spleens of TCR Rg mice analyzed 10wk or 20wk after BM transfer. **(D, E)** TCR transduced NOD.*scid* BM were transferred alone (mono) or co-transferred with NOD.CD45.2 BM (mixed) intravenously into NOD.TCRαKO mice. Recipients were sacrificed 5-7wk after BM transfer. (D) Thymic TCR retrogenic Treg frequencies are plotted against frequencies of CD45.1+ cells in CD4SP thymocytes. (E) Thymic Foxp3+ frequencies within TCR transduced immature CD4SP thymocytes (CD45.1+CD4+CD8-CD3+CD5+CD73-). Data are plotted as means ± SEM (n=3-10). ns > 0.05 by (A&B) one-way ANOVA followed by Benjamini, Krieger and Yekutieli's test (all comparisons are made with 14H4 negative control), (C) multiple Welch t test with Benjamini, Krieger and Yekutieli's correction. (D) Pearson correlation and (E) Welch t test. Outliers were identified and removed from analysis using ROUT method (Q=1).

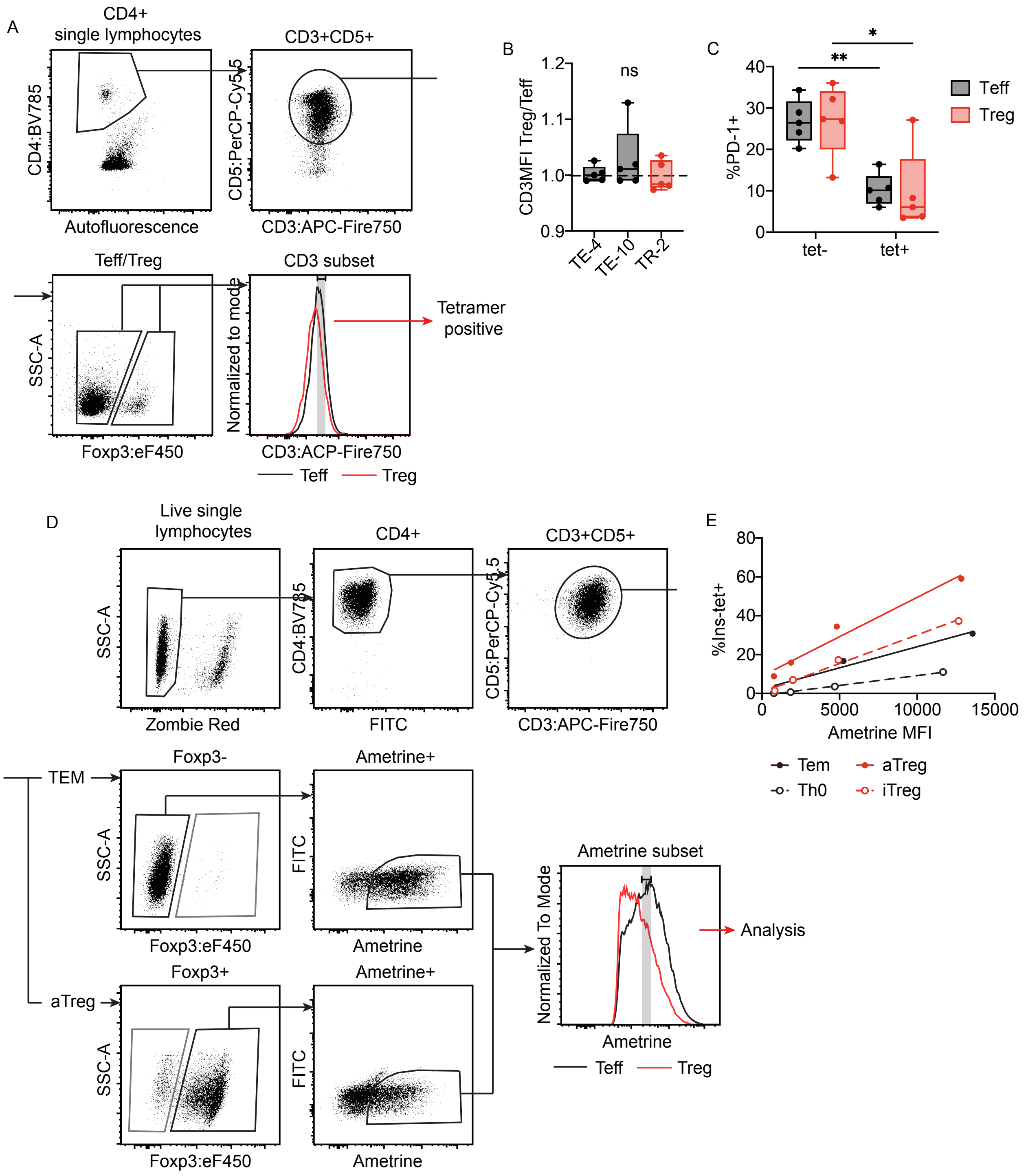


**Supplementary Figure 7. Supplementary data to Figure 5. (A)** Gating strategy for comparing Ins-tet staining on splenic Tregs and Teffs of TCR Rg mice sacrificed 10wk or 20wk after BM transfer. **(B)** Cell surface CD3 expression (Treg versus Teff) after gating for a narrow range of CD3 as demonstrated in (A). Data were analyzed by one-sample t test (n=4-5). **(C)** PD-1 expression on tetramer positive and negative splenic Tregs and Teffs of TR-2 Rg mice. One experiment (n=5). Data analyzed by two-way ANOVA followed by Benjamini, Krieger and Yekutieli's test. **(D)** Gating strategy for comparing Ins-tet staining or Nur77-GFP upregulation in TCR transduced polyclonal NOD Tem and aTreg cells. **(E)** Correlation between Ametrine MFI and Ins-tet+ frequencies. Cells were stained with Ins-tet for one hour on ice. Representative of at least three independent experiments. **(F)** Frequencies of Ins-tet+ cells. Tem and aTreg cells were stained with tetramer for 20min at room temperature followed by 40min on ice. Data are pooled from three independent experiments and analyzed by paired t test. **, p≤0.01; *, p ≤ 0.05; ns > 0.05.

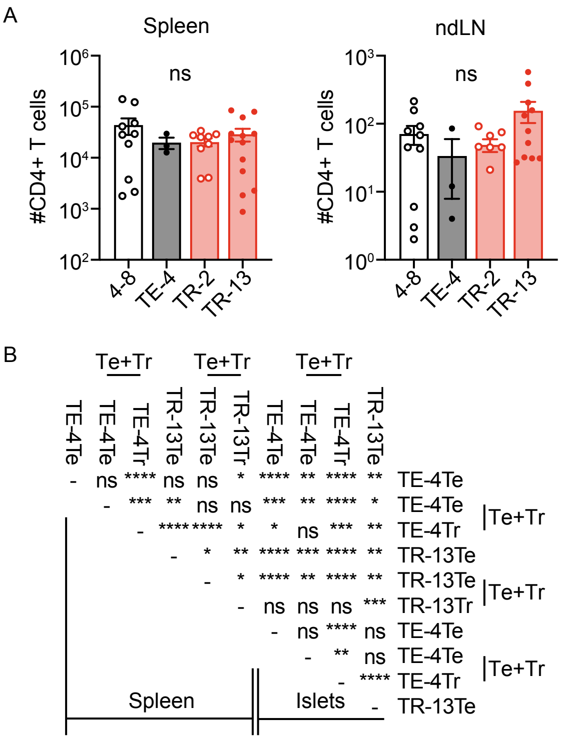


**Supplementary Figure 8. Supplementary data to Figure 6. (A)** Teffs from spleens and ndLNs of TCR Rg mice generated using NOD.*scid* BM were sorted and transferred (IP) into NOD.*scid* female recipients. Recipients were sacrificed for analysis 16wk after transfer or at diabetes onset. Numbers of donor T cells in spleens and ndLNs. **(B)** A complete list for statistical comparison results for Nur77-GFP upregulation of donor Tregs and Teff. The summary box plot is shown in Fig. 7I. Data were pooled from two independent experiments. ****, p ≤ 0.0001; ***, p ≤ 0.001; **, p ≤ 0.01; *, p ≤ 0.05; ns > 0.05 by Kruskal-Wallis test followed by Dunn’s test.

**Supplementary Table I. Summary of insulin specific pancreatic islet Treg and Teff TCRβ repertoire sequencing**

| | Teff | | Treg | | Shared | | %Shared of Teff | | %Shared of Treg | |
|---|---|---|---|---|---|---|---|---|---|---|
| | Amino acid | DNA | Amino acid | DNA | Amino acid | DNA | Amino acid | DNA | Amino acid | DNA |
| P2-TCRα | 358 | 584 | 92 | 118 | 35 | 32 | 9.8% | 5.5% | 38.0% | 27.1% |
| P2-TCRα.Y16A | 210 | 267 | 35 | 41 | 10 | 11 | 4.8% | 4.1% | 28.6% | 26.8% |

Numbers of unique TCRβ sequences, and repertoire overlap between islet-infiltrating Ins-tet+ Tregs and Teffs that develop in P2-TCRα fixed alpha-chain mice, or in insulin epitope mutant P2-TCRα.Y16A mice.

P2-TCRα: Samples were sorted and combined from 38 mice at 10-12wk post BM transfer.

P2-TCRα.Y16A: Islet-infiltrating Ins-tet+ Treg and Teff cells were sorted from 25 splenic CD4+ T cell recipients generated using 7 P2-TCRα.Y16A donors.

**Supplementary Table II. Selected InsB:9-23(E21G/R22E) tetramer positive Treg and Teff TCRs from pancreatic islets of P2-TCRα retrogenic mice.**

| | | CDR3β (aa) | Vβ | Dβ | Jβ | Treg/Teff | Sp Teff | Sp Treg |
|---|---|---|---|---|---|---|---|---|
| Teff TCRs | 4 | CASGGWGGNTLYF | 13-02*01 | 02-01*01 | 02-04*01 | 0.00 | 1 | 0 |
| | 6 | CASRDWGDEQYF | 15-01*01 | 02-01*01 | 02-07*01 | 0.00 | 2 | 0 |
| | 7 | CASSAKTNSDYTF | 13-03*01 | unknown | 01-02*01 | 0.00 | 1 | 0 |
| | 10 | CASSPGQGSEQYF | 05-01*01 | 01-01*01 | 02-07*01 | 0.00 | 1 | 0 |
| | 14 | CASSSGGQGYEQYF | 17-01*01 | 01-01*01 | 02-07*01 | 0.00 | 1 | 0 |
| | 15 | CASSSRRQPYEQYF | 16-01*01 | 01-01*01 | 02-07*01 | 0.00 | 1 | 0 |
| | 16 | CAWKGDRLFF | 31-01*01 | 01-01*01 | 01-04*01 | 0.00 | 2 | 0 |
| | 17 | CAWSLTGGGIEQYF | 31-01*01 | 02-01*01 | 02-07*01 | 0.00 | 0 | 0 |
| | 19 | CTCSADTGGEQYF | 01-01*01 | 01-01*01 | 02-07*01 | 0.00 | 2 | |
| Treg TCRs | 1 | CASADWGGNTLYF | 13-02*01 | 02-01*01 | 02-04*01 | 1.65 | 2 | 2 |
| | 2 | CASGDWGGNTLYF | 13-02*01 | 02-01*01 | 01-03*01 | 1.23 | 3 | 0 |
| | 3 | CASGEAWGGAEQYF | 13-02*01 | 02-01*01 | 02-07*01 | 0.56 | 5 | 5 |
| | 5 | CASGLRDRGDTQYF | 13-02*01 | 01-01*01 | 02-05*01 | 1.50 | 5 | 5 |
| | 8 | CASSLRPRGDSGNTLYF | 26-01*01 | 01-01*01 | 01-03*01 | - | 1 | 1 |
| | 9 | CASSPGDSPLYF | 15-01*01 | 01-01*01 | 01-06*01 | 1.00 | 3 | 3 |
| | 11 | CASSQDGTEVFF | 02-01*01 | 02-01*01 | 01-01*01 | - | 4 | 3 |
| | 13 | CASSQGQGYEQYF | 02-01*01 | 01-01*01 | 02-07*01 | 0.14 | 0 | 0 |
| | 18 | CGASGYNNQAPLF | 20-01*01 | unknown | 01-05*01 | - | 1 | 0 |

Treg/Teff: ratios of templates of Treg verses Teff in islets.

Sp Teff, Sp Treg: Number of mice that harbored the TCR clonotype in splenic Teff or Treg compartment, respectively.
